# Striatal-frontal network activation during voluntary task selection under conditions of monetary reward

**DOI:** 10.1101/434399

**Authors:** Joseph M. Orr, Michael J. Imburgio, Jessica A. Bernard, Marie T. Banich

## Abstract

During voluntary task selection there are a number of internal and external biases that may guide such a choice. However, it is not well understood how reward influences task selection when multiple options are possible. To address this issue, we examined brain activation in a voluntary task-switching paradigm while participants underwent fMRI (n=19). To reinforce the overall goal to choose the tasks randomly, participants were told of a large bonus they would receive at the end of the experiment for making random task choices. We also examined how occasional, random rewards influenced both task performance and brain activation. We hypothesized that these transient rewards would increase the value of the just-performed task and therefore bias participants to choose to repeat the same task on the subsequent trial. Contrary to expectations, transient reward had no consistent behavioral effect on subsequent task choice. Nevertheless, the receipt of such rewards did influence activation in brain regions associated with reward processing as well as those associated with goal-directed control. In addition, reward on a prior trial was found to influence activation during task choice on a subsequent trial, with greater activation in a number of executive function regions as compared to no-reward trials. We posit that both the random presentation of transient rewards and the overall task bonus for random task choices together reinforced the goal to choose the tasks randomly, which in turn influenced activation in both reward-related regions and those regions involved in abstract goal processing.

## Introduction

Executive functioning underlies our ability to effortfully guide behavior towards goals. As executive functions require effort, which is aversive at some level (Kool, McGuire, Rosen, & Botvinick, 2010), motivation is often critical for deciding whether or not to exert effort and exercise executive control (Botvinick & Braver, 2014). Moreover, motivation may actually guide one towards goal-driven behavior. For example, the motivation to enjoy your favorite meal may engage executive processes to arrange for a reservation at the restaurant serving that meal or to organize your behavior so you can cook the meal for yourself.

In fact, there is a growing literature on the relationship between reward-driven motivation and executive function. Motivation to rewards has been suggested to lead to enhanced proactive control that is associated with cognitive stability and sustained goal maintenance, as opposed to more transient, reactive control mechanisms (Braver, 2012; Braver, Gray, & Burgess, 2007). As an example, Locke and Braver (2008) compared performance and BOLD activation during an AX-CPT task under conditions of reward and non-reward, and found that reward was associated with increased sustained activation of prefrontal (PFC) and parietal cortices as well as improved proactive control. This behavioral finding was replicated by Chiew and Braver (2014) as well as by Fröber and Dresibach (2014, 2016a), but only when rewards were contingent on task performance and not random.

Relatedly, Müller and colleagues (2007) examined how the effects of reward that are contingent on a participant’s performance influences the balance between stability and flexibility. In a set-shifting paradigm, they found evidence for increased cognitive stability (i.e., decreased effect of distraction) in blocks of trials where participants could earn money for fast accurate responses, compared to control blocks with no reward. However, the stability-flexibility balance was moderated by perceived effort, with greater stability being observed in participants who reported exerting more effort in response to the rewards. Thus, the effect of reward depends not only on reward contingency, but also effort.

One commonly-used task to examine cognitive flexibility is the task-switching paradigm, in which individuals switch between tasks on a trial-by-trial manner. Typically, in such paradigms, a cue indicates which of the two tasks a participant should perform on any given trial. In fact, several behavioral task-switching studies have now demonstrated that under certain reward structures, flexibility may be increased (Fröber & Dreisbach, 2016b; Kleinsorge & Rinkenauer, 2012; Shen & Chun, 2011). In all these studies, switch costs were reduced when reward value increased from the previous trial (e.g., low reward trial followed by a high reward trial) compared to when reward value remained the same or decreased. In contrast, when prospects of a given reward remain high from one trial to the next, stability is observed with a behavioral benefit on task repeats at the expense of increased switch costs.

Studies that examine reward in standard task switching paradigms assess the influence of reward on the speed of switching between tasks, but are limited in the investigation of overall goal selection and maintenance. That is because in such paradigms the “goal” for the current trial is designated by a cue indicating which task should be performed on that trial. One way to overcome issue is to use a voluntary task switching (VTS) paradigm. In this paradigm, participants are free to choose which of two tasks to perform on every trial, and are typically instructed to choose the tasks equally often, yet in a random order (Arrington & Logan, 2004). Task choices on a given trial are influenced by factors specific to that trial as well as global factors related to overall task performance (Arrington & Logan, 2005; Liefooghe, Demanet, & Vandierendonck, 2010; Mayr & Bell, 2006; Orr, Carp, & Weissman, 2012; Orr & Weissman, 2011; Yeung, 2010). Examining the influence of reward on VTS performance at the local, trial-level as well as the global level might provide insight into the mechanisms by which task choices are made, and thus how overall goals are maintained and executed.

A series of recent behavioral studies examined how task choice on VTS paradigms might be affected by changes in reward magnitude in trials immediately preceding the choice. (Fröber & Dreisbach, 2016b). Four of these studies mixed cued and voluntary task switching, with an additional experiment that used a standard VTS paradigm with only voluntary choice trials. Across the studies, the magnitude of reward (low vs. high) was cued at the beginning of a trial, and varied across trials such that reward magnitude could remain the same from one trial to the next, or increase/decrease in magnitude. Participants were more likely to choose to repeat tasks when a high reward was repeated from the previous trial, whereas a change in reward from the previous trial or a repeated low reward resulted in more frequent switches. Thus, reward could either lead to increased flexibility (more frequent switches) or increased stability (more frequent repeats) depending on the recent reward history. Fröber & Dreisbach interpreted these findings as suggesting that repeated high rewards bias towards exploitation of the current situation whereas the possibility of a reward that is increased in size relative to prior rewards biases towards exploration.

While the study by Fröber & Dreisbach (2016b) demonstrated that task choice on a single trial is influenced by reward on the preceding trial, a recent study by Braem (2017) demonstrated that repeated rewards can bias subsequent task choices for the remainder of the task. In this study, all participants performed a block of trials in which they were explicitly cued as to the task to perform, and hence were directed when to switch tasks, followed by a block of trials in which they themselves voluntarily decided when to switch tasks. The rewards were given after each trial in the cued block, but not in the voluntary block. Presentation of the cued trials was further manipulated as one group of participants received more reward on switch trials than repeat trials, while the other group received more reward for repeat trials. The question of interest was whether the reward contingencies that occurred across the cued block would influence choice in the voluntary block. Participants who received larger rewards after task switches in the cued block made more task switches in the subsequent voluntary choice block than participants who received larger rewards after task repeats in the cued block. This pattern was found despite instructions to make random voluntary task choices, as well as a stipulation that deviating from random task choices would make participants ineligible for a contest for the person with the most points. The results of the study suggest that repeated association of reward with task switches or task repeats can have a lasting effect on task choices throughout an entire block.

The current study aims to build upon the work of Fröber & Dreisbach (2016) and Bream (2017) by examining the effect of rewards on task choices on a trial-by trial level as well as overall task choice (throughout the entire experiment). In the aforementioned studies, reward magnitude varied trial-by-trial, but the frequency of reward was predictable. However, our current understanding of the brain’s reward system circuitry suggests that it appears to be most sensitive to unexpected rewards (Schultz, Dayan, & Montague, 1997). Hence, in the current study, unlike Fröber & Dreisbach (2016) and Braem (2017), reward was presented infrequently and at the end of a trial in an unexpected, random manner. Our hypothesis was that by presenting a reward immediately after performing a given task, the value of that task would be temporarily increased (Schultz, 2006), which in turn would increase the probability that participants would choose that task again on the next trial.

With regards to the brain mechanisms that might be involved in strengthening a task choice in response to reward, there are a number of possibilities. One likely prediction is that the striatum should be involved since it is a critical mechanism in processing the receipt of reward (O’Doherty, 2004; O’Doherty et al., 2004; Schultz et al., 1997; Zald, 2004). Another likely candidate is the PFC cortex which has been shown to be activated when task requirements involve the interaction between rewards, goals, and actions (Rushworth & Behrens, 2008). Moreover, the prefrontal cortex is critical for task switching behavior (Brass, Ullsperger, Knoesche, von Cramon, & Phillips, 2005; Kim, Cilles, Johnson, & Gold, 2012). Nonetheless, it is not clear which region(s) of the prefrontal cortex might be most influenced by reward. Work from Kim and colleagues has suggested that across domains of switching (i.e., stimulus, response, or cognitive set), the inferior frontal junction (IFJ) and posterior parietal cortex are active, suggesting a role of these regions in updating and representing task sets (Kim et al., 2012; Kim, Johnson, Cilles, & Gold, 2011). Some work has shown that activity in the IFJ is sensitive to reward-driven motivational changes (Bahlmann, Aarts, & D’Esposito, 2015; Braver, Paxton, Locke, & Barch, 2009). Thus, one possibility is that reward modulates activity of these regions during task switching. However, these regions have been mainly implicated in cued task switching, and not voluntary task switching.

Hence, another potential region that may be involved is the rostral lateral prefrontal cortex (RLPFC) which helps to maintain an overall abstract goal representation. In the standard VTS paradigm the overarching goal is to guide trial-by-trial task choices so that overall the participant chooses between the two tasks equally often and in a random order (Orr & Banich, 2014). The RLPFC is thought to lie at the apex of a gradient of abstraction within the PFC, with the most anterior regions controlling abstract, domain-general information and more posterior regions controlling domain-specific information like actions (Badre, 2008; O’Reilly, 2010); but see recent work suggesting that the dorsolateral PFC can also show “apex”-like characteristics (Badre & Nee, 2017; Nee & D’Esposito, 2016, 2017). More specifically, the RLPFC is thought to be involved in the high-level processing of goals, subgoals, and integrating information across time and stimulus dimensions (e.g., Charron & Koechlin, 2010; Christoff et al., 2001; Koechlin, Basso, Pietrini, Panzer, & Grafman, 1999; Nee & D’Esposito, 2016). Recently, it has become clear that the RLPFC plays a role in decisionmaking. For example, Kovach and colleagues (2012) demonstrated that patients with RLPFC lesions have difficulty tracking recent trends in reward history. Furthermore, the RLPFC may be critical for considering alternatives to the chosen decision (Boorman, Behrens, Woolrich, & Rushworth, 2009) and exploring unfamiliar options (Daw, O’Doherty, Dayan, Seymour, & Dolan, 2006a). Thus, because the VTS relies on an abstract overall representation of the task goal (i.e., to choose randomly across trials) to guide task choice, reward may influence activity of the RLPFC.

Given these considerations we predicted that the effects of rewards would be observed both immediately after receipt of the reward and also prior to the next task choice. We predicted that the receipt of reward, especially because it was received randomly, would be associated with increased activation in regions of the brain typically associated with processing immediate reward, such, as the ventral striatum and ventral medial PFC (McClure, Laibson, Loewenstein, & Cohen, 2004). The more interesting question is the degree to which such rewards would influence the activation of the IFJ involved in task switching and/or the RLPFC, which is involved in maintaining overall task goals. To the degree that a just received reward influences the maintenance and updating of a specific task set, effects would be expected to be observed on activation of the IFJ. To the degree that reward serves to reinforce the overall higher-order goals (e.g., Charron & Koechlin, 2010), and/or the need to integrate and balance information about the recent reward, that would likely bias to repeating the task choice, with the more long-term goal of a 50/50 distribution of task choice, effects should be observed in RLPFC. These predictions are somewhat speculative as to our knowledge, this is the first imaging study to examine the interactions of reward and voluntary task selection.

## Method

### Participants

Twenty-two young healthy adults were recruited from the University of Colorado Boulder community as part of a larger study of the individual differences on executive function (Orr, Smolker, & Banich, 2015; Reineberg, Andrews-Hanna, Depue, Friedman, & Banich, 2015; Reineberg & Banich, 2016; Smolker, Depue, Reineberg, Orr, & Banich, 2015). The median age was 20 years with a range of 19-27. Seven participants identified as female. We had intended on recruiting 25 participants based on an estimated power analysis but were only able to enroll 22 participants. There were technical problems for two participants and one participant only performed one task throughout the whole experiment, yielding useable data for a total of 19 participants.

### Experimental Paradigm

Participants performed a double-registration variant of a voluntary task switching paradigm (Arrington & Logan, 2005; Orr et al., 2012). On each trial participants were allowed to voluntarily choose between two tasks: one involved making a decision about the numerical size of two digits while the other involved making a decision about the actual physical size of the font in which those two digits were presented. Task choices were made prior to the onset of the task stimuli, which consisted of two numbers; one number was larger than 6 and the other was smaller than 4, and one number was presented in a large font and the other in a small font. Participants were instructed to choose the tasks in a random order. To make this idea concrete, they were given examples of stereotyped task sequences and random sequences. After task performance, reward feedback was presented: on 25% of correct trials the feedback indicated that a 10-cent reward was earned and on the other trials the feedback indicated that 0 cents was earned. Reward was pseudo-randomly presented such that there was an even distribution across the two tasks (within each run). The task was presented using E-Prime 2.10 (https://pstnet.com).

A prototypical trial is presented in Figure 1. The choice cue (i.e., ‘?’, presented in Calibri 24 size font) appeared for 350 ms followed by a fixation cross (presented in Calibri size 24 font) that appeared for 1150 ms. Participants chose the task by pressing a button with their left index or middle finger, with the task-button mapping being counterbalanced across participants. The target digits then appeared for 350 ms, followed by a fixation cross that appeared for 1150 ms. Participants responded to the task with their right index and middle fingers, with the index finger mapped to the bottom number and the middle finger mapped to the top number. The physically large number was presented in Calibri size 32 font and the physically small number was presented in Calibri size 18 font. The numerically large number was randomly selected from the set of 7, 8, and 9, and the numerically small number was randomly selected from the set of 1, 2, and 3. Physical size and numerical size were counterbalanced across trials within each block. A reward feedback stimulus then appeared for 1500 ms, which indicated whether a reward was given or not. A reward was indicated by ¢10¢ (presented in Calibri in size 48 font) and no reward was indicated by ¢00¢ (presented in Calibri size 24 font). Each block consisted of 64 trials, with an equal number of congruent (correct answer is the same for both tasks, e.g., top number numerically and physically large) and incongruent trials (correct answer is different for both tasks, e.g., top number numerically large but physically small, as shown in Fig. 1), with the position of the numerically larger and physically larger digit being counterbalanced. The ITI was jittered between 3000 and 7500 ms according to a decreasing pseudo-exponential distribution that favored short ITIs (i.e., 57 trials at 3000 ms, 4 trials at 4500 ms, 2 trials at 6000 ms, and 1 trial at 7500 ms).

**Figure 1.**
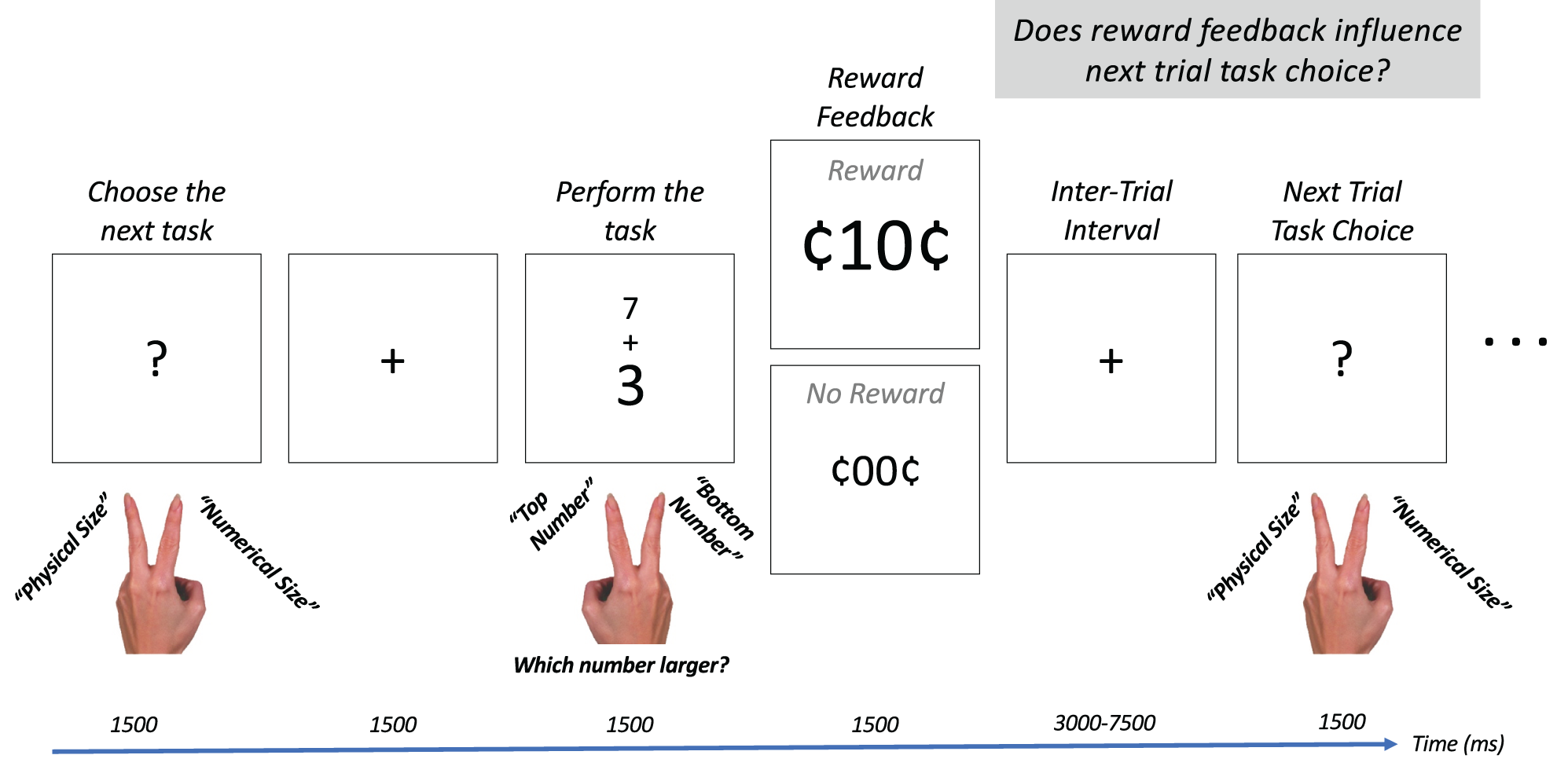
Example trial sequence. At the beginning of each trial participants were cued to choose the next task (either a numerical size comparison or a physical size comparison of two upcoming digits) with a ‘?’. After correct task performance, 25% of trials contained a reward, and the remaining trials did not. We predicted that the reward would increase the value of the just performed task, and would bias participants to choose to perform that task again on the next trial.

Participants were instructed that the trial rewards were random and were used to maintain motivation throughout the experiment. A running total of the rewards was updated at the end of each block. Participants were instructed that the maximum of these trial rewards would be $5. Participants were further instructed that a separate larger bonus, up to $10, would be determined based on how random their task choices were. They were told that the bonus would be delayed by two weeks. In this manner, we were attempting to provide more incentive for random task choices than for task choices biased by the transient rewards.

### fMRI Data Collection

A Siemens Magnetom TIM Trio (3-Tesla) MRI system with a 12-channel head coil was used for data acquisition. Structural images were acquired with a T1-weighted 3D magnetization prepared rapid gradient multi-echo sequence (MPRAGE; sagittal plane; repetition time [TR] = 2530 ms; echo times [TE] = 1.64 ms, 3.5 ms, 5.36 ms, 7.22 ms, 9.08 ms; GRAPPA parallel imaging factor of 2; 1 mm isotropic voxels, 192 interleaved slices; FOV = 256mm; flip angle = 7°; time = 6:03 min). Six functional task runs were collected with an EPI sequence from 26 interleaved slices (3 mm slice thickness, 3.4 × 3.4 mm in-place resolution) aligned parallel to orbital frontal cortex with the following parameters: [TR] = 1500 ms; [TE] = 25 ms; flip angle = 67 deg; 333 measurements; 0.53 ms echo spacing. In order to perform B0 unwarping, a gradient echo fieldmap was collected with the following parameters: TR = 400 ms; TE1 = 4.92 ms; TE2 = 7.38; 3 mm slice with 3.8 mm in-plane resolution. The fieldmap was processed in FSL to generate a phase difference image and a magnitude image for each echo time. B0 unwarping was performed in FEAT as described in more detail below. The structural images were collected in an earlier session occurring an average of 6 months prior.

### fMRI Data Analysis

FMRI data processing was carried out using FEAT (FMRI Expert Analysis Tool) Version 6.00, part of FSL (FMRIB’s Software Library, www.fmrib.ox.ac.uk/fsl). The following prestatistics processing was applied; motion correction using MCFLIRT (Jenkinson, Bannister, Brady, & Smith, 2002); slice-timing correction using Fourier-space time-series phase-shifting; non-brain removal using BET (Smith, 2002); B0 distortion unwarping (Jenkinson, 2003, 2004); spatial smoothing using a Gaussian kernel of FWHM 6.0mm; grand-mean intensity normalisation of the entire 4D dataset by a single multiplicative factor. Preprocessed data were then denoised with ICA-AROMA (Pruim, Mennes, van Rooij, et al., 2015; Pruim, Mennes, Buitelaar, & Beckmann, 2015), an automatic method of removing motion artifacts using ICA implemented with FSL MELODIC. After denoising, the data underwent highpass temporal filtering (Gaussian-weighted least-squares straight line fitting, with sigma=45.0s).

First-level analyses were carried out using FEAT. Time-series statistical analysis was carried out using FILM with local autocorrelation correction (Woolrich, Ripley, Brady, & Smith, 2001). Two models were set-up for each run of the task: 1) BOLD activity associated with the reward feedback stimulus, and 2) BOLD activity associated with task choice as a function of whether participants chose to repeat or switch tasks and whether the previous trial was rewarded or not. Contrasts were defined for the mean of each level (e.g., reward, no reward, repeat, switch), Reward > No Reward, and Switch > Repeat. Thus, there were 3 contrasts for the first model, and 6 for the second model. Registration of the functional data to the T1w and MNI152_T1_2mm template was carried out using FLIRT with a Boundary Based Registration (BBR) cost function (Jenkinson et al., 2002; Jenkinson & Smith, 2001). Registration from the T1w to the MNI152_T1_2mm template was then further refined using FNIRT nonlinear registration (Andersson, Jenkinson, & Smith, 2007a, 2007b).

A second-level model was defined for each participant to average results across the five runs using fixed effect analysis in FEAT. Group-level analyses were performed in *randomize* using non-parametric permutation tests with cluster correction carried out with Threshold-Free Cluster Enhancement (TFCE) (Smith & Nichols, 2009; Winkler, Ridgway, Webster, Smith, & Nichols, 2014) with a cluster-level threshold of FWE-corrected *p* < .05. Note that as the name implies, TFCE does not used a voxel-level threshold and instead relies on a TFCE metric based on the height and width of activation. Some analyses are reported with a higher voxel-level t-statistic threshold in order to yield more separated clusters.

For display purposes, cortical volumetric statistics maps were projected to the HCP 900 Subject Group Average Midthickness Surface (Van Essen et al., 2017) using the Connectome Workbench v. 1.3.0 (https://www.humanconnectome.org/software/get-connectome-workbench), and 2-D slices were created using FSL’s *fsleyes* (v. 0.23.0) on the MNI152_T1_1mm template. Cluster tables were generated using FSL’s automated atlasquery (*autoaq*) using the volumetric data. Cluster labels come primarily from the Harvard-Oxford Cortical and Subcortical atlases, while the Oxford-GSK-Imanova Striatal Connectivity Atlas (Tziortzi et al., 2014) was used for labeling clusters in the striatum, and the Neubert Ventral Frontal Parcellation (Neubert, Mars, Thomas, Sallet, & Rushworth, 2014) and Sallet Dorsal Frontal Parcellation (Sallet et al., 2013) were used for labeling clusters in prefrontal cortex. As the Neubert and Sallet Parcellations were right lateralized, the thresholded t-stat maps were flipped to generate labels for the left hemisphere. Thus, the labels for the left prefrontal clusters may be approximate.

### Regions of Interest

Region of interest (ROI) analyses were conducted in order to facilitate the detection of interactions between previous reward and current task choice, specifically, the decision to repeat or switch. Interactions are difficult to interpret with linear contrasts (e.g., a contrast for the interaction of previous reward and current task alternation is 0 1 −1 0 [Repeat-Reward Repeat-NoReward Switch-Reward Switch-NoReward]). Rather than setting up multiple simple effects contrasts, we chose to conduct a repeated measures ANOVA on Percent Signal Change (PSC) data extracted from *a priori* ROIs in the striatum, IFJ, and RLPFC. Striatal masks were created using the Oxford-GSK-Imanova Striatal Connectivity Atlas, an atlas based on cortical-striatal connectivity (Tziortzi et al., 2014). Striatal masks were created for the executive subregion (connected to areas 9, 9/46 and area 10) and the limbic subregion (connected with orbitofrontal and ventromedial cortex). The atlas masks were thresholded at 60% and 70%, respectively, and binarized. The IFJ ROI was defined from the meta-analysis by Kim and colleagues (2012) and consisted of a 10 mm diameter sphere centered at −40, 3, 33. For the RLPFC ROIs, we focused on the most anterior portion, the frontal pole (FP). Lateral and orbital FP masks were defined from our previous DTI-based connectivity parcellation of the FP (Orr et al., 2015). The FP masks consisted of spheres with a diameter of 10 mm centered on the center of mass from our k=6 parcellation in each hemisphere. Left and right masks were combined into bilateral masks. We previously showed that the lateral FP has more local connections within the PFC than other FP subregions and is functionally connected with the frontoparietal control network. We had previously found that the orbital FP is connected with visual cortex and temporal cortex, including the hippocampus and amygdala.

## Results

### Behavioral Results

#### Task Choice Behavior

We first examined whether the choice to repeat or switch tasks on the next trial (calculated as switch rate, i.e., the proportion of switch trials within a given cell) was influenced by whether or not the previous trial was rewarded. We had predicted that receiving a reward after performing a given task (compared with not receiving a reward) would bias participants to choose to repeat the same task on the next trial. Note that a 50% switch rate indicates a balance between repeating and switching tasks, a switch rate above 50% indicates a bias to switch tasks, and a switch rate below 50% indicates a bias to repeat tasks. Overall, participants showed a preference for switching tasks (M: 53.6% [CI: 50.1-57.3%]; *t*(18) = 2.2, *p* = .043, Cohen’s *d* = 0.50), contrary to most previous studies of voluntary task switching which show a strong repeat bias (e.g., Arrington & Logan, 2004, 2005; Mayr & Bell, 2006; Orr et al., 2012). However, previous studies have shown that even with long (vs. short) response to stimulus intervals, there is a decreased (and sometimes absent) repetition bias (Arrington & Logan, 2004, 2005; Liefooghe, Demanet, & Vandierendonck, 2009; Orr et al., 2012; Yeung, 2010); due to the nature of the slow BOLD response in fMRI, the target to choice stimulus interval was between 4500 and 10500 ms. Whether this long interval explains the switch bias observed here is unclear, and would require behavioral studies with variable intervals. Nevertheless, to our knowledge, no previous study using a standard VTS paradigm has shown a switch bias.

Furthermore, and contrary to our prediction, there was no main effect of whether or not the previous trial was rewarded (No Reward: 52.3% [CI: 48.1% - 56.4%]; Reward: 52.0% [CI: 46.0-58.0%]; F(1,18) = 0.007, *n.s.*, 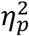 = .000). To further examine this null effect, we used the open source statistical program JASP (JASP Team (2018), Version 0.9 [Computer software]) to calculate the Bayes Factor BF_01_, which quantifies the evidence for the null hypothesis vs. the alternative hypothesis (Rouder, Speckman, Sun, Morey, & Iverson, 2009). This analysis yielded BF_01_ = 4.2, suggesting these data are 4.2 times more likely to be observed under the null hypothesis, reflecting moderate support for accepting the null hypothesis that previous reward had no effect on switch rate.

In contemplating this unexpected outcome, we decided to consider whether the effect of reward on the previous trial affected switch rate based on whether the rewarded trial was a task repeat or task switch trials. We wondered whether participants may have thought that the reward was for repeating or switching tasks, rather than the specific task choice (numerical size, physical size). There was a very large effect of whether or not the previous task had been repeated (Previous Repeat: 60.1%; Previous Switch: 44.8%; F(1,18) = 26.8, *p* < .001, 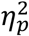 = .60), with individuals more likely to switch tasks on the current trial if they had chosen to repeat on the previous trial (e.g., a task sequence such as Numerical-Numerical-Physical) than if they had chosen to switch on the previous trial (e.g., a task sequence such as Numerical-Physical-Numerical). Critically, however, there was no interaction of previous alteration and previous reward (F(1,18) = 0.24, *n.s.*, 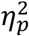 = .013). Thus, it does not appear that participants were guided by the strategy that the transient rewards were associated with the decision to repeat or switch tasks. Nevertheless, with neuroimaging we can look for evidence of reward mechanisms interacting with regions of the brain that underlie the decision to repeat or switch.

#### Task Choice Reaction Time

We considered the effects of task alternation (Repeat, Switch) and previous trial reward (No Reward, Reward) on how quickly a task was chosen (i.e., task choice reaction time). *The results are reported in* Table 1. The speed with which individuals chose a task showed a large effect of alternation (Repeat: 415 ms; Switch: 386 ms; F(1,18) = 15.3, *p* < .001, 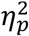 = .46) with participants being faster to choose on trials when they switched task than when they repeated the same tasks. While this finding is inconsistent with previous studies (Arrington & Logan, 2005; Orr & Weissman, 2011), this effect may be related to the switch bias described above. There was a moderate effect on whether the previous trial had been rewarded (No Reward: 392 ms; Reward: 409 ms; F(1,18) = 4.8, *p* = .042, 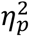 = .21), with participants being slower to choose the next task following a reward as compared to no reward. This suggests that rather than acting on fast heuristics to facilitate task choices, participants may have made more deliberate task choice decisions following a reward.

**Table 1.** Choice reaction time as a function of task alternation and previous trial reward.

| **<u>Task</u>**<br>**<u>Alternation</u>** | **<u>Previous Trial</u>**<br>**<u>Reward</u>** | **<u>Mean</u>** | **<u>SEM</u>** |
| --- | --- | --- | --- |
| Repeat | No Reward | 404 | 23 |
|  | Reward | 426 | 28 |
| Switch | No Reward | 381 | 19 |
|  | Reward | 392 | 24 |

#### Target Performance

Response time and accuracy to the targets (i.e., the digit stimuli) was analyzed as a function of task alternation, *target congruency*, and previous trial reward. *See Table 2 for mean performance*. With regards to reaction time switch costs, participants showed a moderate sized switch cost with RT being longer for trials in which they switched tasks than repeated the same task, but the effect was only at a trend level [Repeat: 497 ms; Switch: 509 ms; F(1,18) = 3.3, *p* = .085, 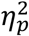 = .16). In terms of accuracy, there was not a significant switch cost (Repeat: 89.6%; Switch: 87.0%; F(1,18) = 2.5, p = .128, 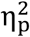 = .12), but there was a significant interaction of task alternation and target congruency (F(1,18) = 4.9, p = .04, 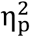 = .16). The switch cost was larger for incongruent (−0.05%) compared to congruent trials (-3.99%), in line with at least one previous study (Orr et al., 2012).

**Table 2.** Target reaction time and accuracy as a function of task alternation, target congruency, and previous trial reward.

| <u>Previous</u><br><u>Trial</u><br><u>Reward</u> | <u>Task</u><br><u>Alternation</u> | <u>Target</u><br><u>Congruency</u> | <u>Reaction Time</u> |  | <u>Accuracy</u> |  |
| --- | --- | --- | --- | --- | --- | --- |
|  |  |  | <u>Mean</u> | <u>SEM</u> | <u>Mean</u> | <u>SEM</u> |
| No Reward | Repeat | Congruent | 465 | 20 | 95.7% | 1.67% |
|  |  | Incongruent | 550 | 26 | 81.6% | 2.84% |
|  | Switch | Congruent | 469 | 19 | 96.0% | 0.99% |
|  |  | Incongruent | 558 | 25 | 77.6% | 2.64% |
| Reward | Repeat | Congruent | 451 | 16 | 98.8% | 0.69% |
|  |  | Incongruent | 534 | 28 | 79.7% | 2.59% |
|  | Switch | Congruent | 463 | 16 | 98.1% | 0.94% |
|  |  | Incongruent | 569 | 40 | 76.2% | 3.33% |

### Whole-Brain Imaging Results

To determine whether indeed our reward manipulation was effective, we first examined brain activation associated with the reward feedback phase of the trial. As expected, the contrast of Reward vs. No Reward feedback yielded significant clusters in the limbic subregion of the striatum, posterior hippocampus, area 47/ frontal operculum (FOp), medial FP, and other regions as reported in Table 3. Figure 2A shows the cortical surfaces while Figure 2B shows striatal slices of the contrast map overlaid on the limbic and executive subregions of the Oxford-GSK-Imanova connectivity striatal atlas (Tziortzi et al., 2014). As shown in Figure 2B, activation after reward feedback was fairly restricted to the limbic subregion of the striatum. Thus, the reward feedback was associated with a number of regions implicated in the receipt of reward as well as internal monitoring of behavior, suggesting that our manipulation was effective in activating reward-related regions of the brain.

**Figure 2.**
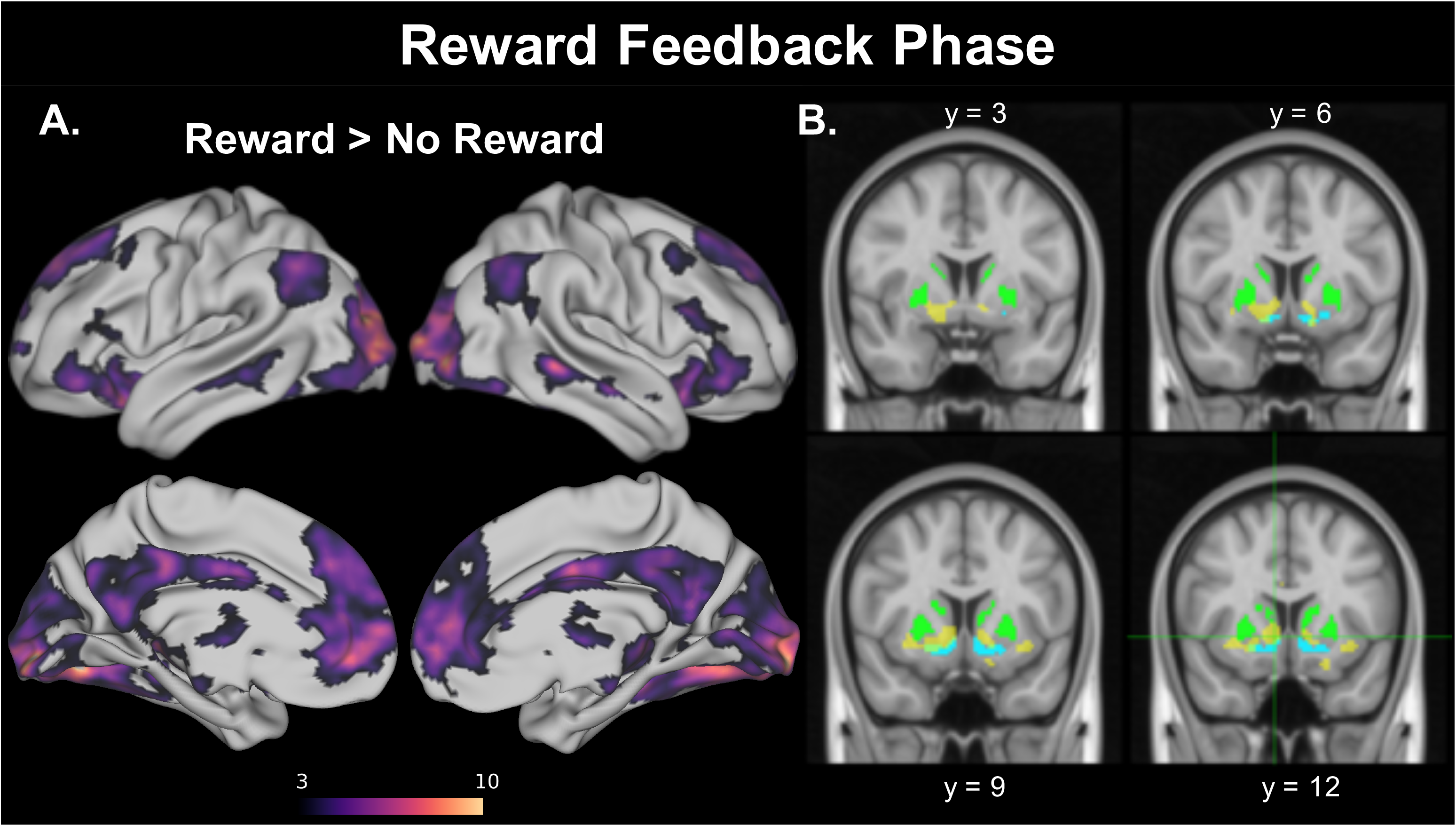
Cortical surface [A] and striatal slices [B] for activation associated with the reward feedback phase. Contrast of reward greater than no reward. Color bar represents t-values with a threshold of t > 5. B. Striatal slices depict activation [yellow] with the striatal executive subregion mask (60% threshold] shown in green and the striatal limbic subregion mask (70% threshold] shown in cyan.

**Table 3.** Atlasquery cluster report from contrast of Reward and No Reward Feedback Stimuli. Additional threshold of t > 5.0 was applied to separate large clusters. Subregions from large clusters are reported below the main table.

| ClusterIndex | Voxels | MAX | MAX X (mm) | MAX Y (mm) | MAX Z (mm) | COG X (mm) | COG Y (mm) | COG Z (mm) | Laterality | Structure label for center of mass |
| --- | --- | --- | --- | --- | --- | --- | --- | --- | --- | --- |
| 26 | 5535 | 12.6 | -26 | -82 | -2 | 1.35 | -84.1 | 2.14 | Bilateral | 37% Intracalcarine Cortex |
| 25 | 1771 | 8.93 | 4 | 66 | 4 | -4.95 | 48.5 | 18.4 | Medial | 77% Paracingulate Gyrus |
| 24 | 493 | 8.31 | 8 | 10 | -4 | 17.2 | 8.97 | -5.41 | Bilateral | 58% Striatum Limbic Subregion |
| 23 | 354 | 10.5 | 22 | -28 | 0 | 25.7 | -30.8 | -6.35 | Right | 49% Hippocampus |
| 22 | 277 | 7.6 | -58 | -64 | 34 | -53.9 | -61.1 | 38 | Left | 58% Lateral Occipital Cortex |
| 21 | 246 | 8.22 | -20 | 14 | -18 | -25.2 | 16.3 | -13.9 | Left | 32% Frontal Orbital Cortex |
| 20 | 173 | 6.5 | -4 | -34 | 36 | 0.827 | -37.3 | 37.5 | Medial | 77% Cingulate Gyrus, posterior division |
| 19 | 147 | 7.02 | 18 | 46 | 50 | 16.6 | 44.4 | 44.9 | Right | 76% Frontal Pole (medial) |
| 18 | 143 | 7.56 | -22 | -36 | -2 | -20.9 | -32.4 | -2.09 | Left | 76% Thalamus |
| 17 | 132 | 7.09 | 4 | -2 | 34 | 1.71 | -5.46 | 32.8 | Medial | 79% Cingulate Gyrus, anterior division |
| 16 | 101 | 7.15 | 42 | 36 | -8 | 40.9 | 35.8 | -9.61 | Right | 52% Neubert Area 47, 44% Neubert Frontal Operculum |
| 15 | 86 | 8.32 | 62 | -38 | -4 | 63 | -35.8 | -4.12 | Right | 35% Middle Temporal Gyrus, posterior division |
| 14 | 59 | 6.34 | -46 | 38 | -10 | -43.4 | 37.7 | -11 | Left | 60% Neubert Area 47 |
| 13 | 52 | 5.96 | 16 | 30 | 52 | 19.3 | 30.9 | 55.2 | Right | 64% Superior Frontal Gyrus |
| 12 | 49 | 6.67 | -34 | 12 | -8 | -32.8 | 13.6 | -7.12 | Left | 42% Insular Cortex |
| 11 | 48 | 6.84 | 68 | -12 | -14 | 66.1 | -11.3 | -13.6 | Right | 64% Middle Temporal Gyrus, posterior division |
| 10 | 38 | 6.7 | 46 | 20 | 16 | 46.8 | 21.2 | 17.6 | Right | 12% Inferior Frontal Gyrus, pars triangularis |
| 9 | 35 | 5.97 | -4 | -52 | 18 | -4.28 | -53.2 | 17.9 | Left | 49% Precuneous Cortex |
| 8 | 32 | 6.1 | 6 | -54 | 18 | 5.42 | -54.1 | 22.6 | Right | 40% Precuneous Cortex |
| 7 | 29 | 6.3 | 0 | 60 | 38 | -1.67 | 58.6 | 37.4 | Medial | Sallet Area 9 |
| 6 | 26 | 5.63 | 44 | -60 | 38 | 44.6 | -60.7 | 40.7 | Right | 41% Lateral Occipital Cortex, superior division |
| 5 | 26 | 6.12 | -6 | -66 | 28 | -5.34 | -66.4 | 29.7 | Left | 55% Precuneous Cortex |
| 4 | 25 | 7.2 | -44 | -46 | -14 | -43.9 | -46.6 | -13.7 | Left | 23% Inferior Temporal Gyrus, temporooccipital part |
| 3 | 25 | 6.07 | 44 | -46 | -14 | 44.7 | -46.1 | -11.8 | Right | 24% Temporal Occipital Fusiform Cortex |
| 2 | 19 | 5.38 | 4 | -70 | 8 | 3.38 | -69.2 | 10.1 | Medial | 53% Intracalcarine Cortex |
| 1 | 17 | 5.49 | -40 | -62 | 46 | -39.5 | -63.2 | 47.1 | Left | 55% Lateral Occipital Cortex, superior division |

Structures to which each cluster belongs to
| Cluster 26 | Average Atlas Probability |
| --- | --- |
| Occipital Pole | 21.0 |
| Occipital Fusiform Gyrus | 10.4 |
| Lateral Occipital Cortex, superior division | 6.7 |
| Lingual Gyrus | 5.7 |
| Lateral Occipital Cortex, inferior division | 3.8 |
| Temporal Occipital Fusiform Cortex | 2.9 |
| Intracalcarine Cortex | 2.3 |
| Cluster 25 | Average Atlas Probability |
| Paracingulate Gyrus | 28.3 |
| Medial Frontal Pole | 22.6 |
| Superior Frontal Gyrus | 8.8 |
| Cingulate Gyrus, anterior division | 5.4 |
| Cluster 22 | Average Atlas Probability |
| Lateral Occipital Cortex, superior division | 32.7 |
| Angular Gyrus | 24.6 |
| Cluster 21 | Average Atlas Probability |
| Neubert Area Frontal Operculum | 44.4 |
| Striatum: Limbic Subregion | 9.7 |

Next, we examined activation for the contrast of switch and repeat trials, time-locked to the choice cue. Compared to repeat trials, switch trials showed greater activation of the Superior Parietal Lobule, Dorsal Premotor Cortex, Frontal Eye Fields, Dorsal Medial Frontal Cortex (including pre-supplementary motor area and cingulate), insula, and thalamus. These results are shown in Figure 3 and reported in Table 4. These regions have been implicated in task switching in a prior fMRI meta-analysis (Kim et al., 2012). It is somewhat surprising that there was no dorsolateral FC activation in this contrast, as this region is frequently associated with task switching and cognitive flexibility, more generally (e.g., Brass & von Cramon, 2002).

**Figure 3.**
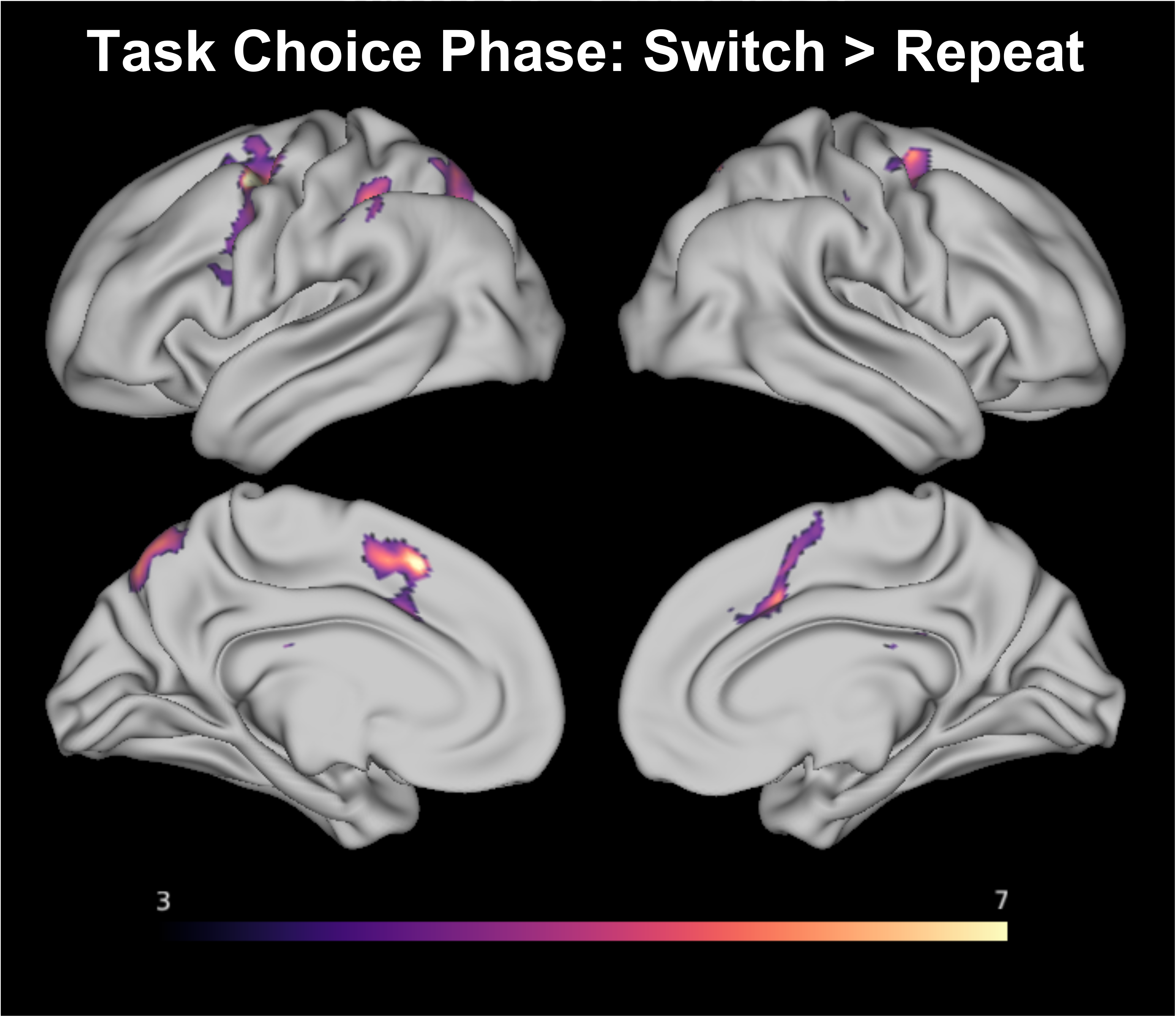
Cortical surface map showing activation of contrast of switch and repeat task choice. Color bar represents t-values with an applied threshold of t > 3.5.

**Table 4.** Atlasquery cluster report from contrast of Switch and Repeat task choices. Additional threshold of t > 3.5 was applied to separate large clusters. Subregions from large clusters are reported below the main table.

| ClusterIndex | Voxels | MAX | MAX X (mm) | MAX Y (mm) | MAX Z (mm) | COG X (mm) | COG Y (mm) | COG Z (mm) | Laterality | Structure label for center of mass |
| --- | --- | --- | --- | --- | --- | --- | --- | --- | --- | --- |
| 10 | 2550 | 7.55 | -38 | -42 | 38 | -28.8 | -53.6 | 45.9 | Left | 38% Superior Parietal Lobule |
| 9 | 1801 | 10.1 | -34 | -4 | 54 | -36.3 | -0.763 | 46.5 | Left | 80% Neubert Area 6v |
| 8 | 1513 | 7.8 | -6 | 20 | 48 | 10.3 | 6.38 | 49.4 | Medial | 44% Juxtapositional Lobule Cortex |
| 7 | 580 | 5.91 | 8 | -30 | 24 | -1.61 | -25.4 | 15.9 | Medial | 70% Left Cerebral White Matter |
| 6 | 330 | 6.32 | 16 | -66 | 56 | 18.8 | -64.4 | 53.7 | Right | 36% Lateral Occipital Cortex |
| 5 | 197 | 5 | 52 | -26 | 42 | 51.1 | -22.8 | 43 | Right | 55% Postcentral Gyrus |
| 4 | 121 | 4.98 | 36 | -36 | 40 | 38.4 | -37.3 | 45.6 | Right | 29% Supramarginal Gyrus |
| 3 | 104 | 4.75 | 44 | 2 | 26 | 47.9 | 3.54 | 31.7 | Right | 54% Precentral Gyrus |
| 2 | 70 | 4.46 | -26 | 10 | 14 | -29.1 | 18 | 10.3 | Left | 100% Neubert 45A, 100% Neubert 44v |
| 1 | 16 | 4.21 | -16 | 4 | 22 | -17.6 | 4.24 | 21.8 | Left | Striatum: 23% Executive Subregion |

Structures to which each cluster belongs to:
| Cluster 10 | Average Atlas Probability |
| --- | --- |
| Lateral Occipital Cortex, superior division | 18.9 |
| Precuneous Cortex | 8.1 |
| Superior Parietal Lobule | 7.9 |
| Supramarginal Gyrus, anterior division | 7.7 |
| Cluster 9 | Average Atlas Probability |
| Sallet Anterior Premotor Dorsal | 16.0 |
| Neubert 6v | 10.8 |
| Neubert IFJ | 8.1 |
| Sallet Area 8A (Frontal Eye Fields) | 5.4 |
| Cluster 8 | Average Atlas Probability |
| Paracingulate Gyrus | 14.7 |
| Juxtapositional Lobule Cortex | 12.4 |
| Cingulate Gyrus, anterior division | 8.9 |
| Cluster 7 | Average Atlas Probability |
| Left Thalamus | 26.0 |
| Left Cerebral White Matter | 25.3 |
| Right Cerebral White Matter | 24.4 |

In the critical analysis, activity during the task choice phase was then examined as a function of the previous trial reward feedback (Reward > No Reward). This contrast yielded a number of clusters including a number of posterior occipital/parietal clusters, posterior dorsolateral PFC (FEF, dorsal premotor), the executive subregion of the striatum, anterior dorsomedial prefrontal cortex (area 9), and RLPFC (lateral FP, 9/46V). Clusters are reported in Table 5, and activation maps are shown in Figure 4. As shown in Figure 4B, the striatal activation is primarily in the executive subregion, in contrast to the effect of reward feedback, which was in the limbic subregion (see Figure 2B). Further comparing Figures 2 and 4, there are several noted changes in regions of activation from receiving reward to choosing the next task following a reward. First, and as already noted, there is a shift from limbic to executive striatum. In the medial PFC, there was a dorsal shift from rostral anterior cingulate/ medial FP to medial area 9. In the lateral prefrontal cortex, there was an anterior shift from area 47/ IFS to the RLPFC (lateral FP/ area 9/46V). In general, this suggests that there was a shift from reward evaluation toward goal processing.

**Table 5.**
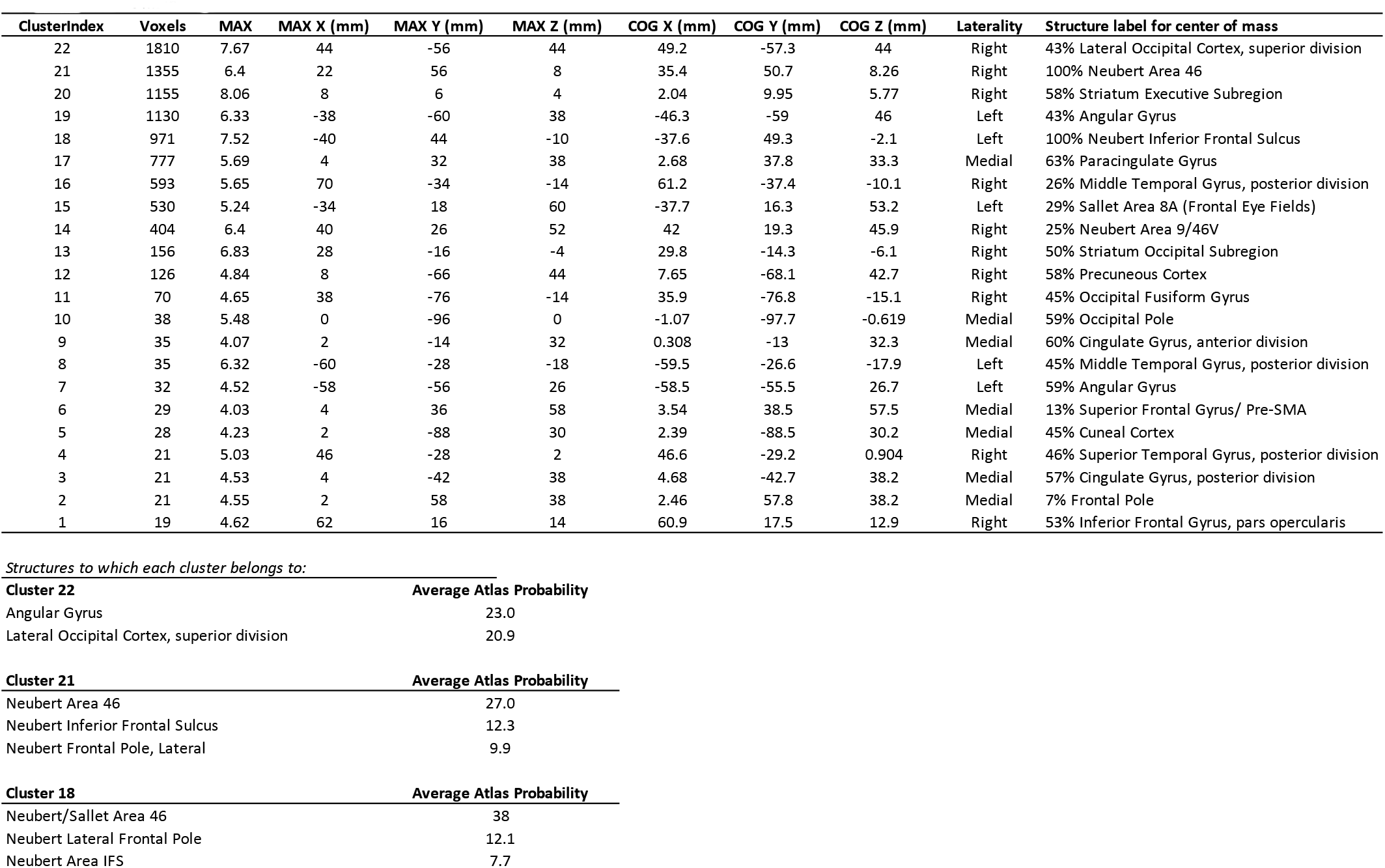
Atlastquery cluster report from contrast of task choices following Reward vs No Reward. Additional threshold of t > 3.5 was applied to separate large clusters. Subregions from large clusters are reported below the main table.

| ClusterIndex | Voxels | MAX | MAX X (mm) | MAX Y (mm) | MAX Z (mm) | COG X (mm) | COG Y (mm) | COG Z (mm) | Laterality | Structure label for center of mass |
| --- | --- | --- | --- | --- | --- | --- | --- | --- | --- | --- |
| 22 | 1810 | 7.67 | 44 | -56 | 44 | 49.2 | -57.3 | 44 | Right | 43% Lateral Occipital Cortex, superior division |
| 21 | 1355 | 6.4 | 22 | 56 | 8 | 35.4 | 50.7 | 8.26 | Right | 100% Neubert Area 46 |
| 20 | 1155 | 8.06 | 8 | 6 | 4 | 2.04 | 9.95 | 5.77 | Right | 58% Striatum Executive Subregion |
| 19 | 1130 | 6.33 | -38 | -60 | 38 | -46.3 | -59 | 46 | Left | 43% Angular Gyrus |
| 18 | 971 | 7.52 | -40 | 44 | -10 | -37.6 | 49.3 | -2.1 | Left | 100% Neubert Inferior Frontal Sulcus |
| 17 | 777 | 5.69 | 4 | 32 | 38 | 2.68 | 37.8 | 33.3 | Medial | 63% Paracingulate Gyrus |
| 16 | 593 | 5.65 | 70 | -34 | -14 | 61.2 | -37.4 | -10.1 | Right | 26% Middle Temporal Gyrus, posterior division |
| 15 | 530 | 5.24 | -34 | 18 | 60 | -37.7 | 16.3 | 53.2 | Left | 29% Sallet Area 8A (Frontal Eye Fields) |
| 14 | 404 | 6.4 | 40 | 26 | 52 | 42 | 19.3 | 45.9 | Right | 25% Neubert Area 9/46V |
| 13 | 156 | 6.83 | 28 | -16 | -4 | 29.8 | -14.3 | -6.1 | Right | 50% Striatum Occipital Subregion |
| 12 | 126 | 4.84 | 8 | -66 | 44 | 7.65 | -68.1 | 42.7 | Right | 58% Precuneus Cortex |
| 11 | 70 | 4.65 | 38 | -76 | -14 | 35.9 | -76.8 | -15.1 | Right | 45% Occipital Fusiform Gyrus |
| 10 | 38 | 5.48 | 0 | -96 | 0 | -1.07 | -97.7 | -0.619 | Medial | 59% Occipital Pole |
| 9 | 35 | 4.07 | 2 | -14 | 32 | 0.308 | -13 | 32.3 | Medial | 60% Cingulate Gyrus, anterior division |
| 8 | 35 | 6.32 | -60 | -28 | -18 | -59.5 | -26.6 | -17.9 | Left | 45% Middle Temporal Gyrus, posterior division |
| 7 | 32 | 4.52 | -58 | -56 | 26 | -58.5 | -55.5 | 26.7 | Left | 59% Angular Gyrus |
| 6 | 29 | 4.03 | 4 | 36 | 58 | 3.54 | 38.5 | 57.5 | Medial | 13% Superior Frontal Gyrus/ Pre-SMA |
| 5 | 28 | 4.23 | 2 | -88 | 30 | 2.39 | -88.5 | 30.2 | Medial | 45% Cuneal Cortex |
| 4 | 21 | 5.03 | 46 | -28 | 2 | 46.6 | -29.2 | 0.904 | Right | 46% Superior Temporal Gyrus, posterior division |
| 3 | 21 | 4.53 | 4 | -42 | 38 | 4.68 | -42.7 | 38.2 | Medial | 57% Cingulate Gyrus, posterior division |
| 2 | 21 | 4.55 | 2 | 58 | 38 | 2.46 | 57.8 | 38.2 | Medial | 7% Frontal Pole |
| 1 | 19 | 4.62 | 62 | 16 | 14 | 60.9 | 17.5 | 12.9 | Right | 53% Inferior Frontal Gyrus, pars opercularis |

|  |  |
| --- | --- |
| Angular Gyrus | 23.0 |
| Lateral Occipital Cortex, superior division | 20.9 |

**Cluster 21** **Average Atlas Probability**
|  |  |
| --- | --- |
| Neubert Area 46 | 27.0 |
| Neubert Inferior Frontal Sulcus | 12.3 |
| Neubert Frontal Pole, Lateral | 9.9 |

**Cluster 18** **Average Atlas Probability**
|  |  |
| --- | --- |
| Neubert/Sallet Area 46 | 38 |
| Neubert Lateral Frontal Pole | 12.1 |
| Neubert Area IFS | 7.7 |

**Figure 4.**
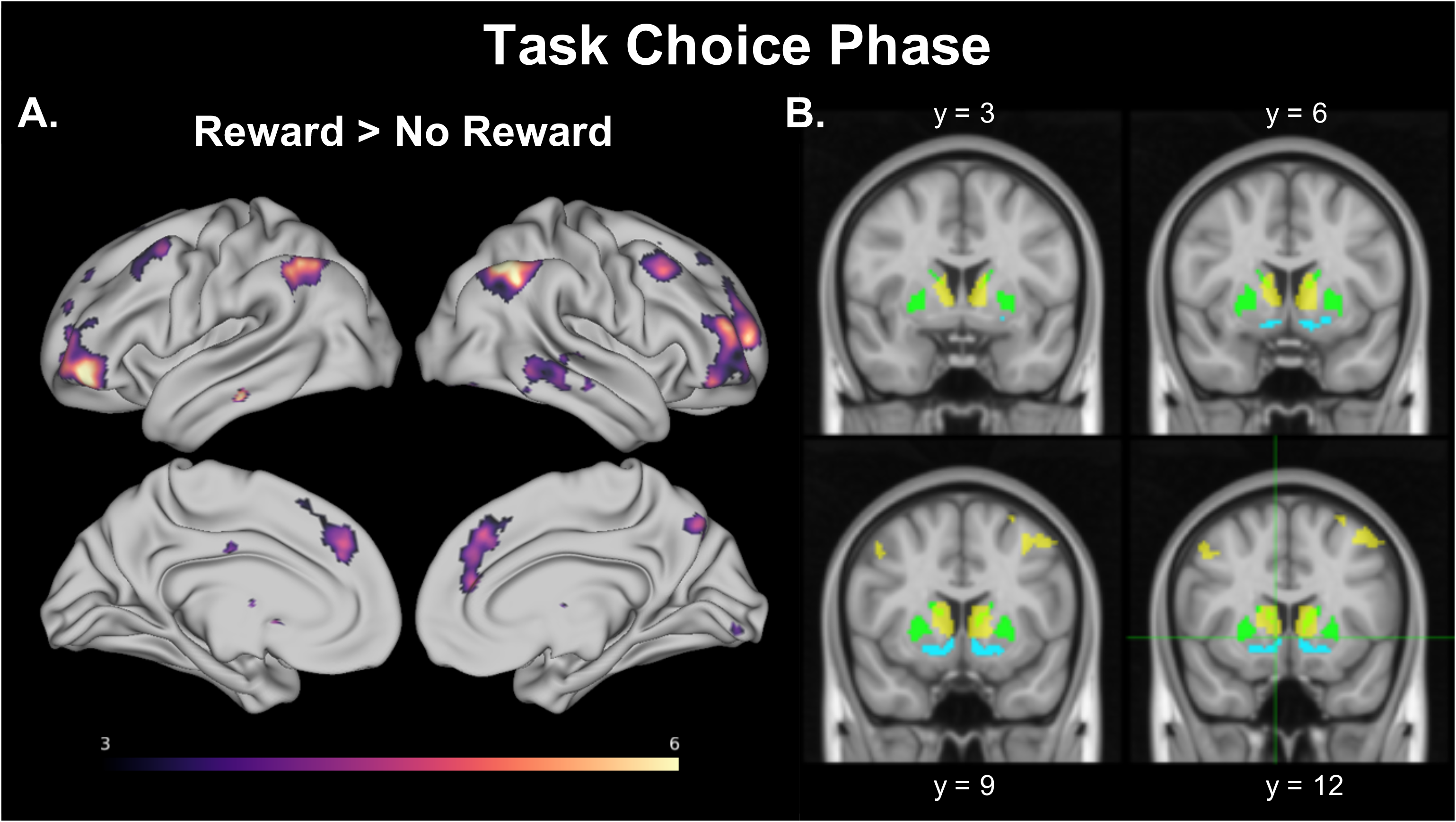
Cortical surface (A) and striatal slices (B) for activation associated with the task choice phase as a function of previous trial reward. Contrast of reward greater than no reward, with the color bar showing a threshold of t>3.5. B. Striatal slices depict activation (yellow) with the striatal executive subregion mask (60% threshold) shown in green and the striatal limbic subregion mask (70% threshold) shown in cyan.

### ROI Analysis

We conducted two separate ROI analyses, one using activation from the reward feedback phase of the trial and another using activation from the task choice phase of the trial. In both cases mean percent signal change (PSC) was extracted from the ROIs. The same five ROIs were used in both: limbic and executive striatal subregions (Tziortzi et al., 2014), IFJ (Kim et al., 2012) and lateral and orbital FP (Orr et al., 2015). The main purpose of the ROI analyses was to test whether reward on the prior trial affected whether to switch vs. repeat tasks on the current trial, but we also looked at the effect of receiving reward feedback. For the analysis of the reward feedback phase, we found a moderate effect of region (F(2.3, 41.6) = 11.1, *p* = 6.8E-5, 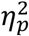 = .38) and a large effect of reward (F(1,18) = 25.1, *p* = 9.1E-5, 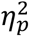 = .58), as well as a moderately-sized interaction effect of region and reward (F(3.1, 55.0) = 4.7, *p* = .005, 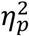 = .22). As shown in Figure 5, simple main effects tests revealed significant effects of reward in all of the ROIs except the IFJ (IFJ: F = 0.12, p = .74; all other ROIs: all F’s > 10.0, all p’s < .002).

**Figure 5.**
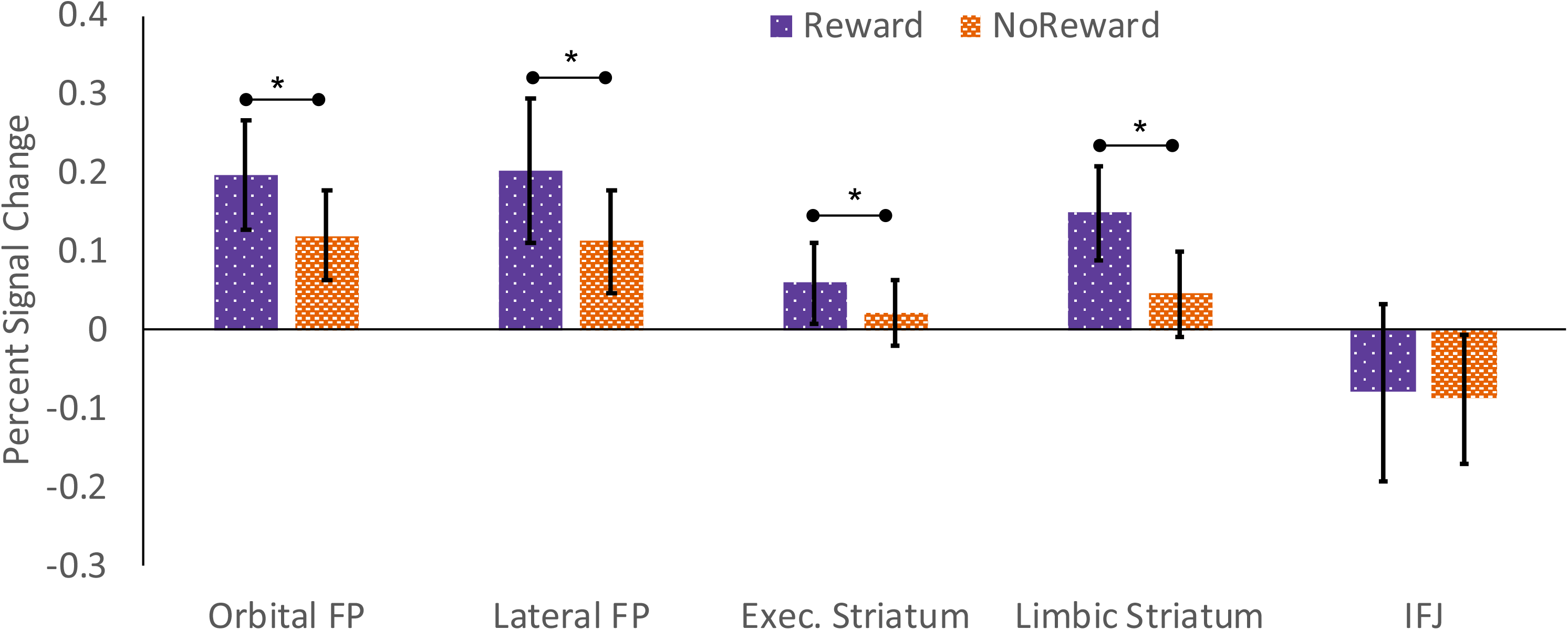
Percent signal change for the reward feedback stimuli. Significant main effects of reward in a given region are denoted by bars and asterisks. FP = Frontal Pole; IFJ = Inferior Frontal Junction.

For the analysis of the task choice phase, there were significant main effects of region (F(2.5, 45.3) = 19.3, *p* = 1.4E-7, 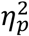 = .52) and reward (F(1, 18) = 4.7, *p* = .04, 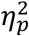 = .21). As shown in Figure 6, PSC was generally greater following reward vs. no reward trials, but showed little effect of whether or not the task choice repeated from the prior trial or was switched. *There was a small, but significant region by previous reward interaction (F(2.7,49.3) = 2.9, p = .28*, 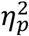 = .*14*), so we examined the simple main effect of reward in the different ROIs. Reward was significant for the Lateral FP (F = 5.9, p = .03) and Executive Striatum (F = 16.7, p = 6.9E-4), but not in the other ROIs (all F’s < 2.6, all p’s > .13). Thus, as predicted, reward-based task choices were associated with the lateral FP and the dorsal striatum. The lack of an effect of task alternation in the IFJ is somewhat surprising, given its critical role in cued task switching (Kim et al., 2012), but it is not clear whether the IFJ is as important in voluntary task switching.

**Figure 6.**
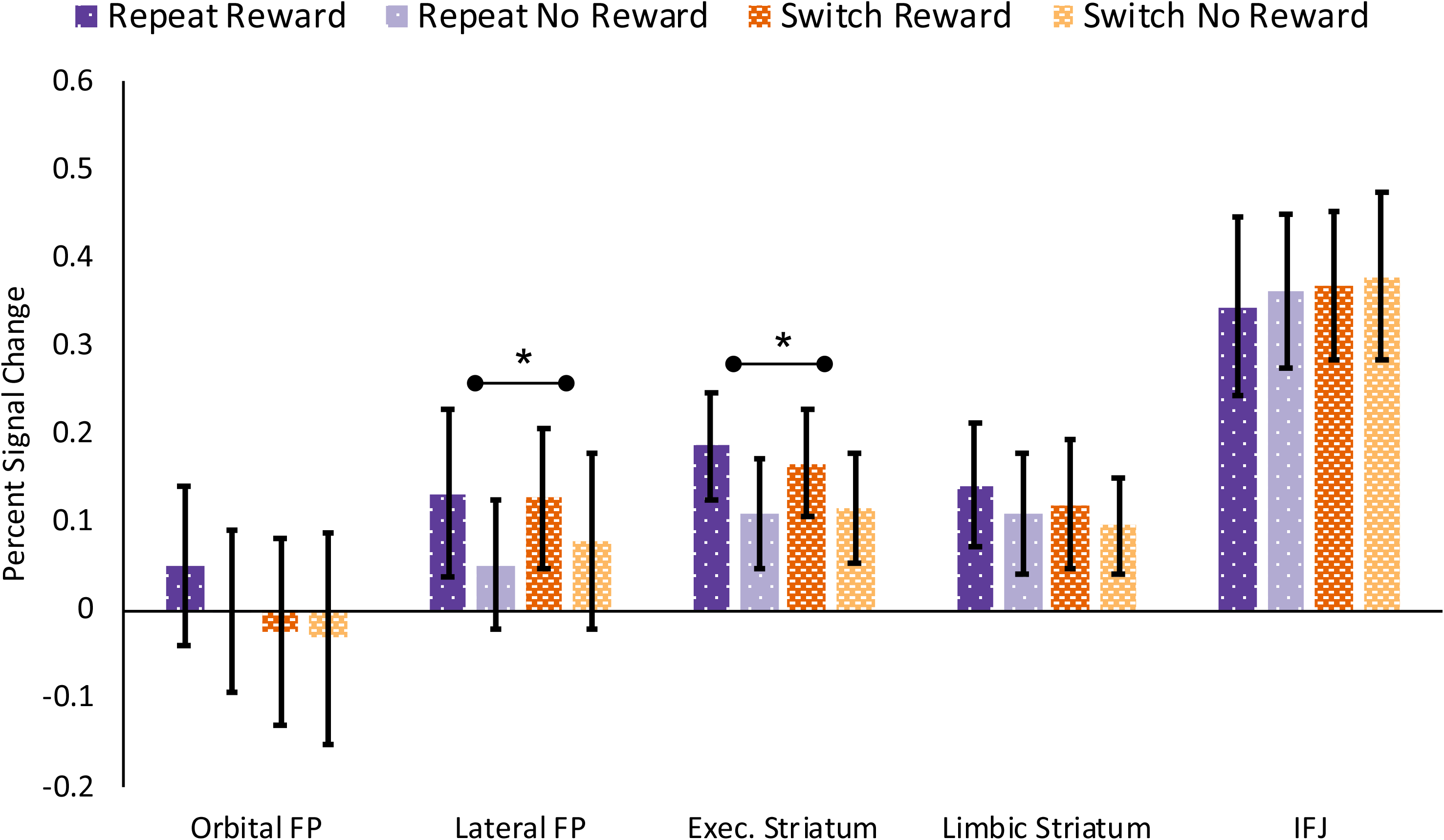
Percent signal change for the task choice period as a function of previous trial reward and current task alternation. Significant main effects of reward in a given region are denoted by bars and asterisks. No significant main effects of alternation were identified. FP = Frontal Pole, IFJ = Inferior Frontal Junction.

## Discussion

In the current study we investigated the influence of reward on behavior and brain activity in a voluntary task switching paradigm. Our results indicated that while activation of striatal regions increased after the receipt of random rewards (that were not contingent on the participant’s performance), such activation did not affect task choice on the subsequent trials. Rather, it appears that the effect of a reward was to influence activation prior to the subsequent task choice in rostrolateral cortex, but not the IFJ, two regions implicated in task choice and task switching, but which have distinct roles. We discuss each of these findings in turn below. First, we discuss the null effects in the behavioral data.

The prediction that *random, non-contingent* rewards would influence task choice was based on an influential literature that demonstrates that the brain’s reward system is sensitive to unexpected rewards (Schultz et al., 1997). However, a growing literature on the role of reward and executive function suggests that the balance between cognitive flexibility vs. stability is influenced by at least two factors: reward-behavior contingencies, and sequential changes in reward magnitude. With regards to reward-behavior contingencies, when rewards are contingent upon behavioral outcomes (i.e., speed cutoffs, accuracy), proactive control appears to be increased, which reinforces stability at the expense of flexibility (Chiew & Braver, 2014; Fröber & Dreisbach, 2016a). On the other hand, rewards that are not contingent on behavior lead to reduced proactive control (Fröber & Dreisbach, 2016a). Relatedly, work on conflict adaptation (i.e., increased reactive control following a high conflict trial vs. a low conflict trial) has further shown that reward feedback presented at the end of the trial, as in the current study, has differential effects on conflict adaptation depending on performance contingencies. When reward is presented randomly without performance contingencies, conflict adaptation is reduced following gains compared to losses or neutral feedback. This occurs presumably because the positive affect from gains counteracts the increased effort required for increased control (van Steenbergen, Band, & Hommel, 2009, 2012). However, when reward is only presented infrequently, and is contingent on performance, conflict adaptation is enhanced following gains (Braem, Verguts, Roggeman, & Notebaert, 2012; Stürmer, Nigbur, Schacht, & Sommer, 2011)]

Sequential changes in reward magnitude, on the other hand, appears to be more closely linked to flexibility. Increased magnitude of potential reward from the previous to the current trial, relative to decreased or consistent reward prospects, has been associated with reduced switch costs and increased proportion of voluntary task switches (Fröber & Dreisbach, 2016b; Kleinsorge & Rinkenauer, 2012; Shen & Chun, 2011). In each of these studies, participants could receive reward for producing responses faster than some criterion (e.g., top percentile of RTs in the previous block). However, prior studies have not examined how voluntary task choice is influenced by the effects of reward that is not contingent on participant performance. Furthermore, in these prior studies, the reward magnitude available on each trial was presented at the beginning of the trial in order to influence task preparation or choice, whereas in the current study, reward feedback was only presented at the end of the trial, after participants had already performed the task. It remains to be seen how non-contingent reward presented at the *beginning* of a trial might influence the stability/flexibility trade-off.

The studies discussed above primarily examined the local effects of a reward, that is, how task performance on the trial immediately following a reward is affected. Recently, Braem (2017) found that global patterns of reward (i.e., varying the relative proportion of task alternations and repetitions that were rewarded) could reinforce abstract control representations such as the proportion of voluntary task repetitions and alternations across a longer run of trials. These effects on voluntary switching emerged even though the reward was administered during an earlier series of cued task repetitions and alternations. This study by Braem raises the possibility that in the current study, participants’ task choice patterns were reinforced by the global reward structure. The fact that the transient rewards were not performance-contingent and were randomly presented may have seemed insignificant in comparison to the abstract goal to choose the tasks randomly, which was further reinforced by the end-of-task bonus for random task choices. Indeed, participants showed a overall switch bias, which is observed when participants are asked to generate random sequences (Baddeley, 1996; Baddeley, Emslie, Kolodny, & Duncan, 1998; Rapoport & Budescu, 1997), suggesting that participants were motivated to select the tasks randomly. Future studies might build on the present work by systematically manipulating a number of variables that may impact the influence of reward on task choice including, whether the reward is contingent or not on an individual’s performance on a task, whether its value is indicated prior or after the task, and frequency of reward.

While the imaging results did not reveal an interaction of previous reward and task choice, there were three interesting shifts of activation in reward-related brain regions from the phase of the trial in which they experienced receipt of reward to the phase in which they determined their task choice on the subsequent trial. First, in the striatum there was a shift in activation from the ventral limbic subregion to the dorsal executive subregion. This finding is consistent with one influential theory of the dorsal-ventral gradient in the striatum is the actor-critic model (O’Doherty et al., 2004; Sutton & Barto, 1998). This model proposes that the ventral striatum uses temporal difference models to predict future reward associated with the current environment, while the dorsal striatum uses this signal to modify action plans to maximize future reward (Haruno, 2004; O’Doherty, 2004; Tricomi, Delgado, & Fiez, 2004). However, we do not have enough evidence in the current study to form a model of what policy or action plan participants employed to guide their task choices. This is an issue that might be fruitfully explore in future studies.

The second region in which a shift was observed was in the medial PFC. During the receipt of reward there was activation of rostral medial PFC, while during the task choice period, the dorsal medial PFC was activated. This shift from reward receipt to task choice appears to reflect a shift from reward evaluation and internal monitoring to planning action intentions. Soon and colleagues (2008; 2013) have shown that activity in the rostral medial PFC precedes the conscious awareness of both a voluntary motor choice and a voluntary task choice, suggesting that this region encodes abstract intentions. In an fMRI meta-analysis, Nakao and colleagues (2012) suggested that the rostral medial PFC is associated with internally-guided decision making, that is decision making according to internal preferences rather than external contingencies. The dorsal medial PFC cluster identified in the current study also overlaps with the region that has been linked to voluntary motor and task decisions across a number of studies (Demanet, de Baene, Arrington, & Brass, 2013; Forstmann, Brass, Koch, & von Cramon, 2005, 2006). Brass and Haggard’s (2008) “What, When, Whether” model of action posits that dorsal medial PFC regions (anterior to the pre-supplementary motor area) are responsible for representing ‘what’ response to make, and ‘whether’ or not to make it. The cluster we identified during the choice period overlapped with regions ascribed to both of these functions. This shift in the location of activation from reward receipt to task choice is also in line with the idea that task choices were driven by the longer-term reward and goal.

Lastly, in the lateral PFC we observed a shift from activation in the lateral orbitofrontal cortex (area 47/12) during reward receipt to the RLPFC during the selection of the task on the subsequent trial. While the medial orbitofrontal cortex has been fairly well characterized, the function of the lateral portion is less clear. Indeed, when entering the coordinates of the center of gravity for the 47/ Fop clusters (left: −44, 38, −10; right: 41, 36, −10) into the meta-analytic tool *neurosynth.org* (Yarkoni, Poldrack, Nichols, Van Essen, & Wager, 2011), there are only a handful of significant associations for the right cluster (most of which are anatomical terms followed by vague associations such as “emotional information”), while the left cluster is associated with terms related to language and semantics. One potential reason for the lack of clarity regarding the function of this region is that the lateral orbitofrontal cortex is prone to signal dropout with many BOLD sequences, especially with older neuroimaging sequences. For this reason, much of what is known about lateral orbitofrontal function comes from studies of non-human primates or lesion studies in humans.

In that regard, Rushworth and colleagues have demonstrated that in both humans and monkeys, the lateral orbitofrontal cortex is critical for linking stimuli to reward (Noonan et al., 2010; Noonan, Chau, Rushworth, & Fellows, 2017; Noonan, Mars, & Rushworth, 2011; Rudebeck et al., 2008; Rushworth, Noonan, Boorman, Walton, & Behrens, 2011). Lesions to the lateral orbitofrontal cortex impact credit assignment, i.e., the ability to learn the value of stimulus options, but no deficits in comparing the value of different options. Other evidence regarding the putative functions of this region comes from Neubert and colleagues (2014, 2015) who compared the connectivity patterns of the PFC in humans and macaques. A posterior ventrolateral region they labelled 47 or 47/12 was found to correspond most closely to macaque area 47/12 (Petrides & Pandya, 2002), and was functionally coupled to anterior PFC and anterior temporal cortex. They suggested that this region’s connectivity supports its role in credit assignment, allowing it to learn which sensory features in the environment are rewarding.

The role of RLPFC, the region whose activation we observed during task choice, has been suggested to play a distinct role with regards to reward processing from these other frontal regions. For example, Boorman and colleagues (2009) had participants freely choose between two possible response options which were associated with random amounts of reward. However, the reward probabilities depended on recent choice history. Using a Bayesian reinforcement learning model, they showed that the RLPFC tracked the advantage of choosing the alternative action choice, but did not reflect the value of the chosen option. Conversely, they found that the rostral medial PFC reflected the value of the chosen option, but did not encode the alternative action choice. This finding of the rostral medial PFC encoding current value is in line with the current finding that this region was active during the reward feedback phase, but not during the subsequent trial task choice. It should be noted that in Boorman and colleagues’ study, the activation was 10-14 mm more lateral than the lateral frontal pole ROI used here, and corresponds to area 9/46. Future research is needed to test whether the RLPFC activity observed in the current study during task choice reflects a consideration of choosing the alternative task choice.

However, most previous studies have focused on medial-lateral functional gradients within the RLPFC, not dorsal-ventral. In our previous parcellation of the RLPFC, we found evidence for a medial-lateral gradient as well as a dorsal-ventral gradient (Orr et al., 2015). We suggested that the medial areas of RLPFC are involved in monitoring and regulation of behavior and emotion while lateral areas maintain abstract cognitive representations; we posited that ventral RLPFC is involved in linking stimuli to values and emotions while dorsal RLPFC is involved in abstract action planning. The ROI analysis in the present study demonstrated that the orbital FP ROI (ventral lateral portion of the RLPFC) was involved during the receipt of reward but was not active during the task choice period. The involvement of the orbital FP during the receipt of reward is in line with the suggestion that ventral portions of the RLPFC are involved with linking stimuli to values. The more dorsal lateral FP ROI however, was active during the task choice period, which is in line with the suggestion that lateral dorsal RLPFC is involved with maintaining abstract cognitive representations and action plans. Future studies should further explore the functional differentiation of the frontal pole subregions.

While we found an effect of reward on activation of RLPFC, we did not do so for the IFJ. We thought that IFJ might show such effect for two reasons. First, it is an area that has been implicated in task switching. For example, Forstmann and colleagues (2005) have found that IFG is involved in updating tasks according to both abstract cues to repeat or switch tasks as well as explicit task cues. In addition, there is evidence that the IFJ might be influenced by reward. For example, Bahlmann and colleagues (2015) found that functional coupling between the dopaminergic midbrain and IFJ was correlated with increased cognitive flexibility in a cued switching paradigm, suggesting that reward motivation enhances dopaminergic projections to the IFJ if flexible updating of task information is needed. In the current study, however, we found no influence of reward on activity in the IFJ. While Forstmann and colleagues (2005) found that the IFJ is involved in both internally-guided and explicit task switches, previous imaging work with the voluntary task switching paradigm has not identified a critical role for the IFJ (Demanet et al., 2013; Forstmann et al., 2006; Orr & Banich, 2014). Although the IFJ ROI did not show an effect of alternation, the whole brain contrast of switch > repeat trials identified a posterior frontal cluster that was partially located in IFJ, though primarily in dorsal premotor cortex. (see Figure 3 and Table 4). Thus, it appears that the IFJ does not play a role in voluntary task switching.

### Limitations and Future Directions

While the results of our study are interesting, the study is not without limitations. It is well known that individuals vary in their sensitivity to reward (Braver, 2012; Corr, 2004; Depue & Collins, 1999; Gray, 1970). Yet the relatively low number of participants (N=19) in our study precluded the analysis of individual differences. Such analyses might have allowed for finer-grained analysis of how such individual differences might influence the trade-off between reward-induced biases and the goal to be random and/or the pattern of brain activation associated with each.

Another limitation is that our task design did not allow us to directly compare the reward feedback phase and task choice phase results. Rather our approach was to model each separately in two different GLM analyses. If one wanted to more explicitly examine shifts in brain activation from the feedback phase to the task choice phase, the use of a Finite Impulse Response model would allow one to model the BOLD timeseries. However, given the variable delay between reward and next trial task choice, such an analysis was not feasible for the current experiment, but might be employed with a modified design in future studies.

Finally, one might want to expand on the current study to examine the influence of reward structure in a paradigm that juxtaposes exploration as compared to exploitation of task choice, such as that used by Braun and Arrington (2017). It would be interesting to see whether the same brain structures implicated in our study are also involved in mediating between these two strategies. Indeed previous work has implicated the RLPFC in supporting exploration strategies and the striatum and rostral medial PFC for exploitative strategies (Daw, O’Doherty, Dayan, Seymour, & Dolan, 2006b).

## Conclusions

While we predicted that participants would show a task choice bias induced by the transient rewards, there was no reliable effect of reward on the next trial’s task choice. Nevertheless, these rewards were associated with significant activation of reward-related brain regions. Task choices following the reward showed increased activation of RLPFC regions thought to be involved in high-level abstract control. Interestingly, the effect of reward prior to task selection on the subsequent trial was limited to RLPFC and was not observed for IFJ. Combined with previous work showing that reward can reinforce abstract control representation, these findings suggest that task choices were guided by abstract goals to choose the tasks randomly as implemented by anterior PFC regions. Overall, this study adds to a growing literature on the role of RLPFC in executive function and decision-making. Moreover, this study represents an initial step towards understanding how reward influences voluntary task selection.

## References

Andersson, J. L. R., Jenkinson, M., & Smith, S. (2007a). Non-linear registration, aka spatial normalisation. FMRIB Technial Report TR07JA2.

Andersson, J. L. R., Jenkinson, M., & Smith, S. M. (2007b). Non-linear optimisation. FMRIB technical report TR07JA1. Retrieved from http://fsl.fmrib.ox.ac.uk/analysis/techrep/tr07ja1/tr07ja1.pdf

Arrington, C. M., & Logan, G. D. (2004). The cost of a voluntary task switch. Psychological Science, 15(9), 610–5. https://doi.org/10.1111/j.0956-7976.2004.00728.x

Arrington, C. M., & Logan, G. D. (2005). Voluntary task switching: chasing the elusive homunculus. Journal of Experimental Psychology: Learning, Memory, and Cognition, 31(4), 683–702. https://doi.org/10.1037/0278-7393.31.4.683

Baddeley, A. D. (1996). Exploring the central executive. Quarterly Journal of Experimental Psychology, 49A(1), 5–28. https://doi.org/10.1080/713755608

Baddeley, A. D., Emslie, H., Kolodny, J., & Duncan, J. (1998). Random generation and the executive control of working memory. Quarterly Journal of Experimental Psychology, 51A(4), 819–852. https://doi.org/10.1080/027249898391413

Badre, D. (2008). Cognitive control, hierarchy, and the rostro-caudal organization of the frontal lobes. Trends in Cognitive Sciences, 12(5), 193–200. https://doi.org/10.1016/j.tics.2008.02.004

Badre, D., & Nee, D. E. (2017). Frontal Cortex and the Hierarchical Control of Behavior. Trends in Cognitive Sciences, 22(2), 170–188. https://doi.org/10.1016/j.tics.2017.11.005

Bahlmann, J., Aarts, E., & D’Esposito, M. (2015). Influence of Motivation on Control Hierarchy in the Human Frontal Cortex. Journal of Neuroscience, 35(7), 3207–3217. https://doi.org/10.1523/JNEUR0SCI.2389-14.2015

Boorman, E. D., Behrens, T. E. J., Woolrich, M. W., & Rushworth, M. F. S. (2009). How green is the grass on the other side? Frontopolar cortex and the evidence in favor of alternative courses of action. Neuron, 62(5), 733–743. https://doi.org/10.1016/j.neuron.2009.05.014

Botvinick, M. M., & Braver, T. S. (2014). Motivation and Cognitive Control: From Behavior to Neural Mechanism. Annual Review of Psychology, (September 2014), 1–31. https://doi.org/10.1146/annurev-psych-010814-015044

Braem, S. (2017). Conditioning task switching behavior. Cognition, 166, 272–276. https://doi.org/10.1016/j.cognition.2017.05.037

Braem, S., Verguts, T., Roggeman, C., & Notebaert, W. (2012). Reward modulates adaptations to conflict. Cognition, 125(2), 324–332. https://doi.org/10.1016/j.cognition.2012.07.015

Brass, M., & Haggard, P. (2008). The what, when, whether model of intentional action. The Neuroscientist, 14(4), 319–25. https://doi.org/10.1177/1073858408317417

Brass, M., Ullsperger, M., Knoesche, T. R., von Cramon, D. Y., & Phillips, N. a. (2005). Who comes first? The role of the prefrontal and parietal cortex in cognitive control. Journal of Cognitive Neuroscience, 17(9), 1367–75. https://doi.org/10.1162/0898929054985400

Brass, M., & von Cramon, D. Y. (2002). The role of the frontal cortex in task preparation. Cerebral Cortex, 12(9), 908–14. Retrieved from http://www.ncbi.nlm.nih.gov/pubmed/12183390

Braver, T. S. (2012). The variable nature of cognitive control: a dual mechanisms framework. Trends in Cognitive Sciences, 16(2), 106–13. https://doi.org/10.1016/j.tics.2011.12.010

Braver, T. S., Gray, J. R., & Burgess, G. C. (2007). Explaining the many varieties of working memory variation: Dual mechanisms of cognitive control. In C. Jarrold (Ed.), Variation in Working Memory (pp. 76–106). Oxford: Oxford University Press. https://doi.org/10.3758/s13423-011-0165-y

Braver, T. S., Paxton, J. L., Locke, H. S., & Barch, D. M. (2009). Flexible neural mechanisms of cognitive control within human prefrontal cortex. Proceedings of the National Academy of Sciences of the United States of America, 106(18), 7351–6. https://doi.org/10.1073/pnas.0808187106

Charron, S., & Koechlin, E. (2010). Divided representation of concurrent goals in the human frontal lobes. Science, 328(5976), 360–363. https://doi.org/10.1126/science.1183614

Chiew, K. S., & Braver, T. S. (2014). Dissociable influences of reward motivation and positive emotion on cognitive control. Cognitive, Affective and Behavioral Neuroscience, 14(2), 509–529. https://doi.org/10.3758/s13415-014-0280-0

Christoff, K., Prabhakaran, V., Dorfman, J., Zhao, Z., Kroger, J. K., Holyoak, K. J., & Gabrieli, J. D. (2001). Rostrolateral prefrontal cortex involvement in relational integration during reasoning. NeuroImage, 14(5), 1136–1149. https://doi.org/10.1006/nimg.2001.0922

Corr, P. J. (2004). Reinforcement sensitivity theory and personality. Neuroscience & Biobehavioral Reviews, 28(3), 317–332.

Daw, N. D., O’Doherty, J. P., Dayan, P., Seymour, B., & Dolan, R. J. (2006a). Cortical substrates for exploratory decisions in humans. Nature, 441(7095), 876–879. https://doi.org/10.1038/nature04766

Daw, N. D., O’Doherty, J. P., Dayan, P., Seymour, B., & Dolan, R. J. (2006b). Cortical substrates for exploratory decisions in humans. Nature, 441(7095), 876–879. https://doi.org/10.1038/nature04766

Demanet, J., de Baene, W., Arrington, C. M., & Brass, M. (2013). Biasing free choices: The role of the rostral cingulate zone in intentional control. NeuroImage, 72, 207–213. https://doi.org/10.1016/j.neuroimage.2013.01.052

Depue, R. A., & Collins, P. F. (1999). Neurobiology of the structure of personality: Dopamine, facilitation of incentive motivation, and extraversion. Behavioral and Brain Sciences, 22(03). https://doi.org/10.1017/S0140525X99002046

Forstmann, B. U., Brass, M., Koch, I., & von Cramon, D. Y. (2005). Internally generated and directly cued task sets: an investigation with fMRI. Neuropsychologia, 43(6), 943–52. https://doi.org/10.1016/j.neuropsychologia.2004.08.008

Forstmann, B. U., Brass, M., Koch, I., & von Cramon, D. Y. (2006). Voluntary selection of task sets revealed by functional magnetic resonance imaging. Journal of Cognitive Neuroscience, 18(3), 388–98. https://doi.org/10.1162/089892906775990589

Fröber, K., & Dreisbach, G. (2014). The differential influences of positive affect, random reward, and performance-contingent reward on cognitive control. Cognitive, Affective and Behavioral Neuroscience, 14(2), 530–547. https://doi.org/10.3758/s13415-014-0259-x

Fröber, K., & Dreisbach, G. (2016a). How performance (non-)contingent reward modulates cognitive control. Acta Psychologica, 168, 65–77. https://doi.org/10.1016/j.actpsy.2016.04.008

Fröber, K., & Dreisbach, G. (2016b). How sequential changes in reward magnitude modulate cognitive flexibility: Evidence from voluntary task switching. Journal of Experimental Psychology: Learning Memory and Cognition, 42(2), 285–295. https://doi.org/10.1037/xlm0000166

Gray, J. A. (1970). The psychophysiological basis of introversion-extraversion. Behaviour Research and Therapy, 8(3), 249–266. https://doi.org/10.1016/0005-7967(70)90069-0

Haruno, M. (2004). A Neural Correlate of Reward-Based Behavioral Learning in Caudate Nucleus: A Functional Magnetic Resonance Imaging Study of a Stochastic Decision Task. Journal of Neuroscience, 24(7), 1660–1665. https://doi.org/10.1523/JNEUROSCI.3417-03.2004

Jenkinson, M. (2003). Fast, automated,N-dimensional phase-unwrapping algorithm. Magnetic Resonance in Medicine, 49(1), 193–197. https://doi.org/10.1002/mrm.10354

Jenkinson, M. (2004). Improving the registration of B0-disorted EPI images using calculated cost function weights. NeuroImage, 22, e1544–e1545.

Jenkinson, M., Bannister, P., Brady, M., & Smith, S. M. (2002). Improved Optimization for the Robust and Accurate Linear Registration and Motion Correction of Brain Images. NeuroImage, 17(2), 825–841. https://doi.org/10.1006/nimg.2002.1132

Jenkinson, M., & Smith, S. M. (2001). A global optimisation method for robust affine registration of brain images. Medical Image Analysis, 5(2), 143–156. Retrieved from http://www.ncbi.nlm.nih.gov/pubmed/11516708

Kim, C., Cilles, S. E., Johnson, N. F., & Gold, B. T. (2012). Domain general and domain preferential brain regions associated with different types of task switching: A Meta-Analysis. Human Brain Mapping, 33(1), 130–142. https://doi.org/10.1002/hbm.21199

Kim, C., Johnson, N. F., Cilles, S. E., & Gold, B. T. (2011). Common and distinct mechanisms of cognitive flexibility in prefrontal cortex. Journal of Neuroscience, 31(13), 4771–9. https://doi.org/10.1523/JNEUROSCI.5923-10.2011

Kleinsorge, T., & Rinkenauer, G. (2012). Effects of monetary incentives on task switching. Experimental Psychology, 59(4), 216–226. https://doi.org/10.1027/1618-3169/a000146

Koechlin, E., Basso, G., Pietrini, P., Panzer, S., & Grafman, J. (1999). The role of the anterior prefrontal cortex in human cognition. Nature, 399(6732), 148–151. https://doi.org/10.1038/20178

Kool, W., McGuire, J. T., Rosen, Z. B., & Botvinick, M. M. (2010). Decision Making and the Avoidance of Cognitive Demand. Journal of Experimental Psychology: General, 139(4), 665–682. https://doi.org/10.1037/a0020198

Kovach, C. K., Daw, N. D., Rudrauf, D., Tranel, D., O’Doherty, J. P., & Adolphs, R. (2012). Anterior prefrontal cortex contributes to action selection through tracking of recent reward trends. Journal of Neuroscience, 32(25), 8434–42. https://doi.org/10.1523/JNEUROSCI.5468-11.2012

Liefooghe, B., Demanet, J., & Vandierendonck, A. (2009). Is advance reconfiguration in voluntary task switching affected by the design employed? Quarterly Journal of Experimental Psychology, 62(5), 850–7. https://doi.org/10.1080/17470210802570994

Liefooghe, B., Demanet, J., & Vandierendonck, A. (2010). Persisting activation in voluntary task switching: it all depends on the instructions. Psychonomic Bulletin & Review, 17(3), 381–6. https://doi.org/10.3758/PBR.17.3.381

Locke, H. S., & Braver, T. S. (2008). Motivational influences on cognitive control: Behavior, brain activation, and individual differences. Cognitive, Affective, & Behavioral Neuroscience, 8(1), 99–112. https://doi.org/10.3758/CABN.8.1.99

Mayr, U., & Bell, T. (2006). On how to be unpredictable: evidence from the voluntary task-switching paradigm. Psychological Science, 17(9), 774–80. https://doi.org/10.1111/j.1467-9280.2006.01781.x

McClure, S. M., Laibson, D. I., Loewenstein, G., & Cohen, J. D. (2004). Separate neural systems value immediate and delayed monetary rewards. Science, 306(5695), 503–507. https://doi.org/10.1126/science.1100907

Müller, J., Dreisbach, G., Goschke, T., Hensch, T., Lesch, K. P., & Brocke, B. (2007). Dopamine and cognitive control: The prospect of monetary gains influences the balance between flexibility and stability in a set-shifting paradigm. European Journal of Neuroscience, 26(12), 3661–3668. https://doi.org/10.1111/j.1460-9568.2007.05949.x

Nakao, T., Ohira, H., & Northoff, G. (2012). Distinction between externally vs. Internally guided decision-making: Operational differences, meta-analytical comparisons and their theoretical implications. Frontiers in Neuroscience, 6(MAR), 1–26. https://doi.org/10.3389/fnins.2012.00031

Nee, D. E., & D’Esposito, M. (2016). The hierarchical organization of the lateral prefrontal cortex. ELife, 5, 1–26. https://doi.org/10.7554/eLife.12112

Nee, D. E., & D’Esposito, M. (2017). Causal evidence for lateral prefrontal cortex dynamics supporting cognitive control. ELife, 6. https://doi.org/10.7554/eLife.28040

Neubert, F.-X., Mars, R. B., Sallet, J., & Rushworth, M. F. S. (2015). Connectivity reveals relationship of brain areas for reward-guided learning and decision making in human and monkey frontal cortex. Proceedings of the National Academy of Sciences, 112(20), E2695–E2704. https://doi.org/10.1073/pnas.1410767112

Neubert, F.-X., Mars, R. B., Thomas, A. G., Sallet, J., & Rushworth, M. F. S. (2014). Comparison of human ventral frontal cortex areas for cognitive control and language with areas in monkey frontal cortex. Neuron, 81(3), 700–13. https://doi.org/10.1016/j.neuron.2013.11.012

Noonan, M. P., Chau, B. K. H., Rushworth, M. F. S., & Fellows, L. K. (2017). Contrasting Effects of Medial and Lateral Orbitofrontal Cortex Lesions on Credit Assignment and Decision-Making in Humans. Journal of Neuroscience, 37(29), 7023–7035. https://doi.org/10.1523/JNEUROSCI.0692-17.2017

Noonan, M. P., Mars, R. B., & Rushworth, M. F. S. (2011). Distinct Roles of Three Frontal Cortical Areas in Reward-Guided Behavior. Journal of Neuroscience, 31(40), 14399–14412. https://doi.org/10.1523/JNEUROSCI.6456-10.2011

Noonan, M. P., Walton, M. E., Behrens, T. E. J., Sallet, J., Buckley, M. J., & Rushworth, M. F. S. (2010). Separate value comparison and learning mechanisms in macaque medial and lateral orbitofrontal cortex. Proceedings of the National Academy of Sciences, 107(47), 20547–20552. https://doi.org/10.1073/pnas.1012246107

O’Doherty, J. P. (2004). Reward representations and reward-related learning in the human brain: Insights from neuroimaging. Current Opinion in Neurobiology, 14(6), 769–776. https://doi.org/10.1016/j.conb.2004.10.016

O’Doherty, J. P., Dayan, P., Schultz, J., Deichmann, R., Friston, K. J., & Dolan, R. J. (2004). Dissociable Role of Ventral and Dorsal Striatum in Instrumental Conditioning. Science, 304(5669), 452–454. https://doi.org/10.1126/science.1094285

O’Reilly, R. C. (2010). The What and How of prefrontal cortical organization. Trends in Neurosciences, 33(8), 355–361. https://doi.org/10.1016/j.tins.2010.05.002

Orr, J. M., & Banich, M. T. (2014). The neural mechanisms underlying internally and externally guided task selection. NeuroImage, 84, 191–205. https://doi.org/10.1016/j.neuroimage.2013.08.047

Orr, J. M., Carp, J., & Weissman, D. H. (2012). The influence of response conflict on voluntary task switching: a novel test of the conflict monitoring model. Psychological Research, 76(1), 60–73. https://doi.org/10.1007/s00426-011-0324-9

Orr, J. M., Smolker, H. R., & Banich, M. T. (2015). Organization of the human frontal pole revealed by large-scale DTI-based connectivity: Implications for control of behavior. PLoS One, 10(5). https://doi.org/10.1371/journal.pone.0124797

Orr, J. M., & Weissman, D. H. (2011). Succumbing to bottom-up biases on task choice predicts increased switch costs in the voluntary task switching paradigm. Frontiers in Psychology, 2(February), 31. https://doi.org/10.3389/fpsyg.2011.00031

Petrides, M., & Pandya, D. N. (2002). Comparitive cytoarchitectonic analysis of the human and the macaque ventrolateral prefrontal cortex and the corticocortical connection patterns in the monkey. European Journal of Neuroscience, 16, 291–310. https://doi.org/10.1046/j.1460-9568.2002.02090.x

Pruim, R. H. R., Mennes, M., Buitelaar, J. K., & Beckmann, C. F. (2015). Evaluation of ICA-AROMA and alternative strategies for motion artifact removal in resting state fMRI. NeuroImage, 112, 278–287. https://doi.org/10.1016/j.neuroimage.2015.02.063

Pruim, R. H. R., Mennes, M., van Rooij, D., Llera, A., Buitelaar, J. K., & Beckmann, C. F. (2015). ICA-AROMA: A robust ICA-based strategy for removing motion artifacts from fMRI data. NeuroImage, 112, 267–277. https://doi.org/10.1016/j.neuroimage.2015.02.064

Rapoport, A., & Budescu, D. V. (1997). Randomization in individual choice behavior. Psychological Review, 104(3), 603–617. https://doi.org/10.1037//0033-295X.104.3.603

Reineberg, A. E., Andrews-Hanna, J. R., Depue, B. E., Friedman, N. P., & Banich, M. T. (2015). Resting-state Networks Predict Individual Differences in Common and Specific Aspects of Executive Function. NeuroImage, 104, 69–78. https://doi.org/10.1016/j.neuroimage.2014.09.045

Reineberg, A. E., & Banich, M. T. (2016). Functional connectivity at rest is sensitive to individual differences in executive function: A network analysis. Human Brain Mapping, 37(8), 2959–2975. https://doi.org/10.1002/hbm.23219

Rouder, J. N., Speckman, P. L., Sun, D., Morey, R. D., & Iverson, G. (2009). Bayesian t tests for accepting and rejecting the null hypothesis. Psychonomic Bulletin and Review, 16(2), 225–237. https://doi.org/10.3758/PBR.16.2.225

Rudebeck, P. H., Behrens, T. E. J., Kennerley, S. W., Baxter, M. G., Buckley, M. J., Walton, M. E., & Rushworth, M. F. S. (2008). Frontal Cortex Subregions Play Distinct Roles in Choices between Actions and Stimuli. Journal of Neuroscience, 28(51), 13775–13785. https://doi.org/10.1523/JNEUROSCI.3541-08.2008

Rushworth, M. F. S., & Behrens, T. E. J. (2008). Choice, uncertainty and value in prefrontal and cingulate cortex. Nature Neuroscience, 11(4), 389–97. https://doi.org/10.1038/nn2066

Rushworth, M. F. S., Noonan, M. P., Boorman, E. D., Walton, M. E., & Behrens, T. E. J. (2011). Frontal Cortex and Reward-Guided Learning and Decision-Making. Neuron, 70(6), 1054–1069. https://doi.org/10.1016/j.neuron.2011.05.014

Sallet, J., Mars, R. B., Noonan, M. P., Neubert, F.-X., Jbabdi, S., O’Reilly, J. X.,…Rushworth, M. F. S. (2013). The organization of dorsal frontal cortex in humans and macaques. Journal of Neuroscience, 33(30), 12255–12574. https://doi.org/10.1523/JNEUROSCI.5108-12.2013

Schultz, W. (2006). Behavioral Theories and the Neurophysiology of Reward. Annual Review of Psychology, 57(1), 87–115. https://doi.org/10.1146/annurev.psych.56.091103.070229

Schultz, W., Dayan, P., & Montague, P. R. (1997). A Neural Substrate of Prediction and Reward. Science, 275(5306), 1593–1599. https://doi.org/10.1126/science.275.5306.1593

Shen, Y. J., & Chun, M. M. (2011). Increases in rewards promote flexible behavior. Attention, Perception, and Psychophysics, 73(3), 938–952. https://doi.org/10.3758/s13414-010-0065-7

Smith, S. M. (2002). Fast robust automated brain extraction. Human Brain Mapping, 17(3), 143–155. https://doi.org/10.1002/hbm.10062

Smith, S. M., & Nichols, T. E. (2009). Threshold-free cluster enhancement: addressing problems of smoothing, threshold dependence and localisation in cluster inference. NeuroImage, 44(1), 83–98. https://doi.org/10.1016/j.neuroimage.2008.03.061

Smolker, H. R., Depue, B. E., Reineberg, A. E., Orr, J. M., & Banich, M. T. (2015). Individual differences in regional prefrontal grey matter morphometry and fractional anisotropy are associated with different constructs of executive function. Brain Structure & Function, 220(3), 1291–1306. https://doi.org/10.1007/s00429-014-0723-y

Soon, C. S., Brass, M., Heinze, H.-J., & Haynes, J.-D. (2008). Unconscious determinants of free decisions in the human brain. Nature Neuroscience, 11(5), 543–545. https://doi.org/10.1038/nn.2112

Soon, C. S., He, H. H., Bode, S., & Haynes, J.-D. (2013). Predicting free choices for abstract intentions. Proceedings of the National Academy of Sciences of the United States of America, 110(15), 6217–6222. https://doi.org/10.1073/pnas.1212218110

Stürmer, B., Nigbur, R., Schacht, A., & Sommer, W. (2011). Reward and punishment effects on error processing and conflict control. Frontiers in Psychology, 2(November), 335. https://doi.org/10.3389/fpsyg.2011.00335

Sutton, R. S., & Barto, A. C. (1998). Reinforcement Learning: An Introduction. Cambridge, MA: The MIT Press.

Tricomi, E. M., Delgado, M. R., & Fiez, J. A. (2004). Modulation of Caudate Activity by Action Contingency. Neuron, 41(2), 281–292. https://doi.org/10.1016/S0896-6273(03)00848-1

Tziortzi, A. C., Haber, S. N., Searle, G. E., Tsoumpas, C., Long, C. J., Shotbolt, P.,…Gunn, R. N. (2014). Connectivity-Based Functional Analysis of Dopamine Release in the Striatum Using Diffusion-Weighted MRI and Positron Emission Tomography. Cerebral Cortex, 24(5), 1165–1177. https://doi.org/10.1093/cercor/bhs397

Van Essen, D. C., Smith, J., Glasser, M. F., Elam, J., Donahue, C. J., Dierker, D. L.,…Harwell, J. (2017). The Brain Analysis Library of Spatial maps and Atlases (BALSA) database. NeuroImage, 144, 270–274. https://doi.org/10.1016/j.neuroimage.2016.04.002

van Steenbergen, H., Band, G. P. H., & Hommel, B. (2009). Reward counteracts conflict adaptation. Evidence for a role of affect in executive control. Psychological Science, 20(12), 1473–7. https://doi.org/10.1111/j.1467-9280.2009.02470.x

van Steenbergen, H., Band, G. P. H., & Hommel, B. (2012). Reward valence modulates conflict-driven attentional adaptation: Electrophysiological evidence. Biological Psychology, 90(3), 234–241. https://doi.org/10.1016/j.biopsycho.2012.03.018

Winkler, A. M., Ridgway, G. R., Webster, M. a, Smith, S. M., & Nichols, T. E. (2014). Permutation Inference for the General Linear Model. NeuroImage, 92, 381–97. https://doi.org/10.1016/j.neuroimage.2014.01.060

Woolrich, M. W., Ripley, B. D., Brady, M., & Smith, S. M. (2001). Temporal Autocorrelation in Univariate Linear Modeling of FMRI Data. NeuroImage, 14(6), 1370–1386. https://doi.org/10.1006/nimg.2001.0931

Yarkoni, T., Poldrack, R. A., Nichols, T. E., Van Essen, D. C., & Wager, T. D. (2011). Large-scale automated synthesis of human functional neuroimaging data. Nature Methods, 8(8), 665–675. https://doi.org/10.1038/NMETH.1635

Yeung, N. (2010). Bottom-up influences on voluntary task switching: the elusive homunculus escapes. Journal of Experimental Psychology: Learning, Memory, and Cognition, 36(2), 348–62. https://doi.org/10.1037/a0017894

Zald, D. H. (2004). Dopamine Transmission in the Human Striatum during Monetary Reward Tasks. Journal of Neuroscience, 24(17), 4105–4112. https://doi.org/10.1523/JNEUROSCI.4643-03.2004

